# Sensor Location Optimization and Joint Angle Prediction in Hand Postures

**DOI:** 10.1101/157784

**Authors:** Nayan Bhatt, SKM Varadhan

**Affiliations:** Department of Applied Mechanics, Indian Institute of Technology Madras, Chennai, India

## 1 Sensor selection problem

Grasping of objects (Mason et al. 2001; Santello et al. 2002) or hand shaping during grasping is of interest in robotics, neuroscience, and biomechanics. In animation industry and biomechanics, tracking body and hand movements are essential. For tracking body movements and hand movements, a sizable number of markers are required. As number of marker increase complexity and cost also increase. Researchers have tried various approaches for selecting a set of minimum no. of markers for accurate reproduction of movements. Data-driven approaches have been proposed for classification of various grasps. Some studies have focussed on prediction of joint angles (Kang et al. 2012; Hoyet et al. 2012; Wheatland et al. 2013). Our objective is to predict joint angles from a reduced set of sensors. For preliminary sorting purpose, we used PCA based sensor ranking which is discussed further.

**Figure 1:**
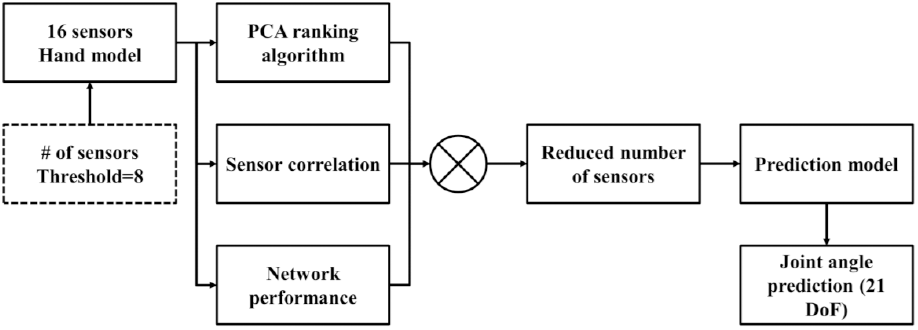
Block diagram representation for sensor selection and joint angle prediction approach. The general block diagram for sensor selection and joint angle prediction is shown in figure 1 above. It shows 16 sensors need to be reduced to the predefined threshold value of 8. Three different algorithms were used for ranking the sensors. Our systematic approach combines these algorithms and gives possibly best reduced set of sensors. These reduced set of sensors were used for predicting all 21 joint angles. All the blocks are discussed in detail in future sections and subsections.

### 1.1 PCA ranking based approach

For the purpose of analysis and ranking, all static postures were considered. Raw data from the sensors, the orientations (yaw, pitch, and roll) are used for computing importance of sensors and ranking all the sixteen sensors. First, we performed PCA on the dataset with 35000 samples and 48 dimensions. Performing PCA on dataset will generate principal component coefficients (i.e. ***W***=48×48) having 48 eigenvectors and corresponding eigenvalues (i.e. λs). Sensor importance for each orientation for each sensor is calculated separately based on multiplying eigenvalues with coefficients of eigenvector (λ_i_ *W_i_).

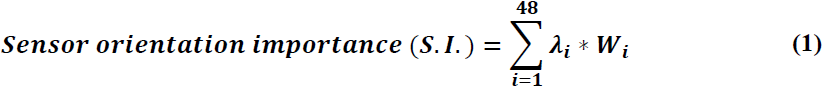

Finally, overall sensor importance is calculated by summing sensor orientation importance across orientations for each sensor.

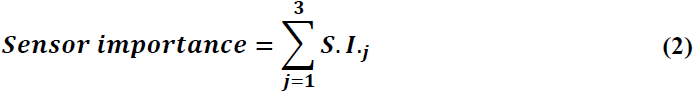

Ranking of the sensor is calculated for each subject separately. All the sensors are arranged in ascending order of *Sensor importance* (i.e. least important sensor on left and most important sensor on the right-hand side) and sensor importance curve is generated. For deciding cutoff on sensors, we took the first derivative of the sensor importance curve and found the peak value. The right-hand side of peak represent sensors that are important from a data point of view.

For the purpose of illustration, consider a single subject static data. The orientation of sensors was used as input matrix to PCA analysis. Size of data matrix is 35000 samples (i.e. 70 points ×5 trials ×100 postures) × 48 variables (16 sensors × 3 orientations). Singular value decomposition algorithm was used for PCA analysis. Analysis generates loading coefficient matrix ***W*** (48 × 48) (i.e. projection matrix), columns represents principal components. Latent represents eigenvalues of eigenvectors (i.e. 48 × 1). Absolute values of coefficient present in eigenvector of ***W*** is multiplied with respective eigenvalues (i.e. λ) and summed up across all the eigenvectors. *Sensor orientation importance* (S.I.) is a vector (i.e. λ*W* = 48 × 1). Matrix *W* and eigenvalues are shown in figure 2 and 3 respectively.

**Figure 2:**
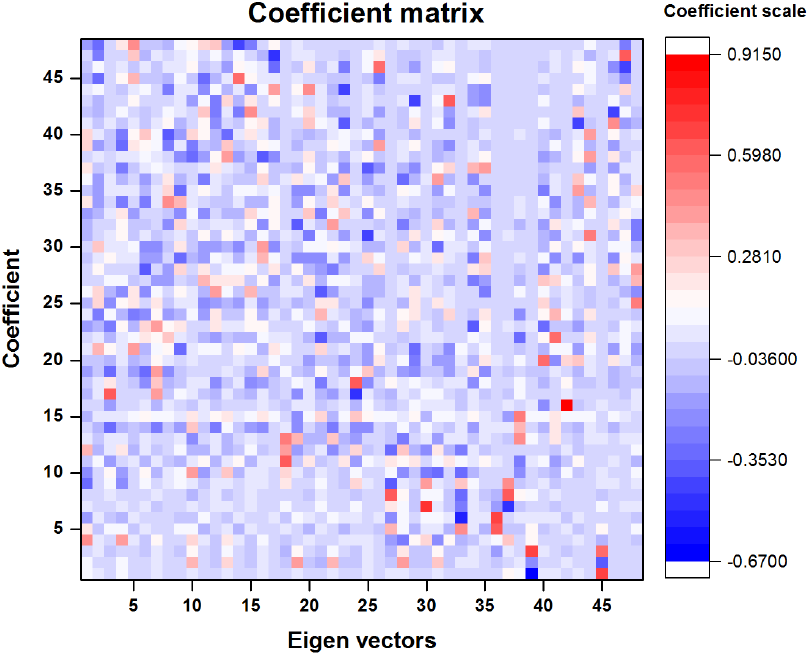
Eigenvector matrix. Columns correspond to eigenvectors. Eigenvectors are ordered in descending order of explained variance.

**Figure 3:**
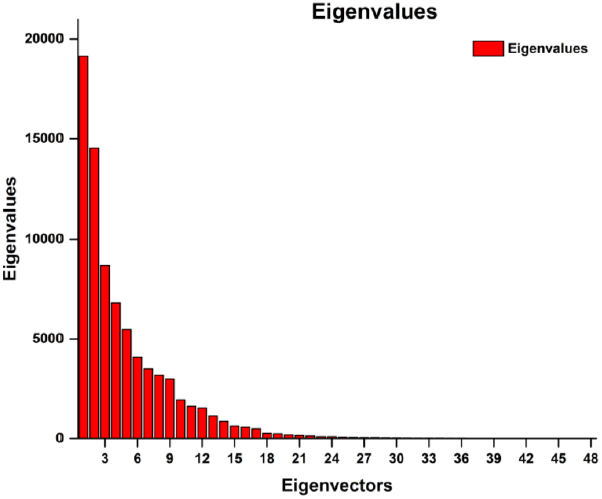
Eigenvalue bar graph for a single subject data. These eigenvalues are used as weights for multiplying coefficient with coefficient matrix W.

Columns of the coefficient matrix represent eigenvectors in descending order of variance. The first column represents first PC representing largest variance in the dataset. The absolute value of coefficient determines the magnitude of loading which is used for ranking the sensors.

Eigenvalues for corresponding eigenvector (i.e. projection W) matrix determines the magnitude of eigenvector in each PC direction. Hence, weights are given to each coefficient based on its position in loading coefficient matrix.

Finally, for computing sensor importance, values of S.I. are summed up for all orientation of figure 4. PCA based rank algorithm gives overall sensor importance as shown in figure 5.

**Figure 4:**
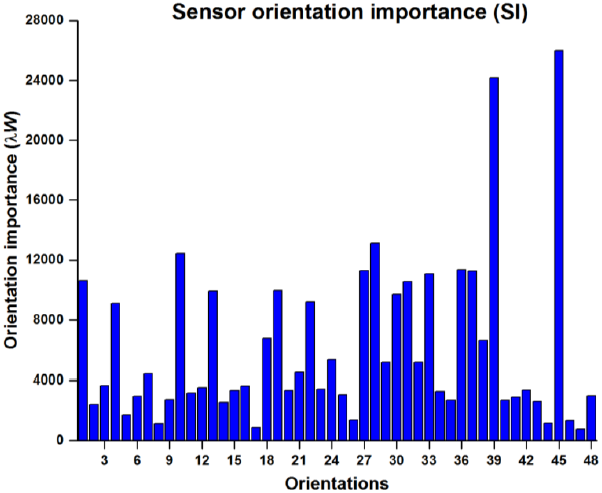
Sensor orientation importance. Each column represents sensor importance calculated based on absolute values of coefficient multiplied with corresponding eigenvalues.

**Figure 5:**
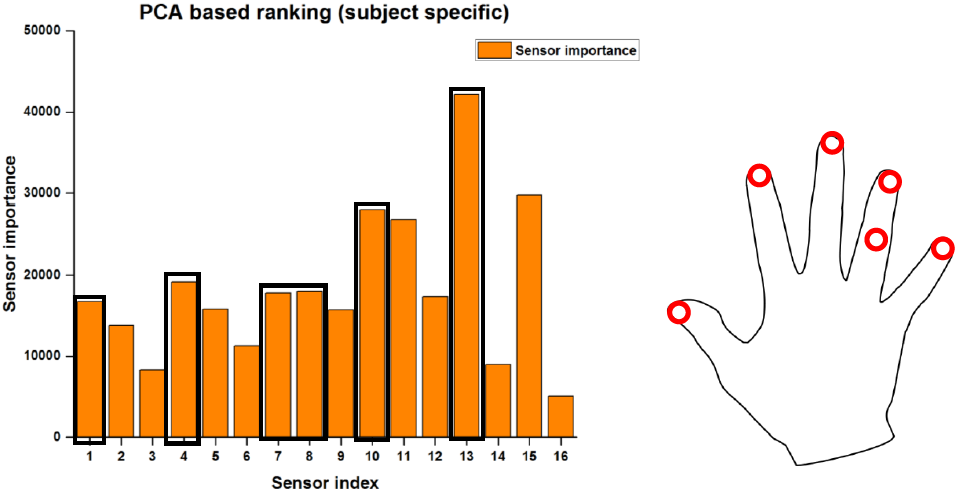
**(a)** PCA-based sensor ranking. Sensor 1, 4, 7, 10, 13 shows relatively large value of sensor rank for a finger (I, M, R, L, and Thumb) for subject 1. Sensor 16 shows the least importance since variance in the wrist is least, gives relatively less importance during PCA analysis. **(b)** Sensor location based on importance calculated from PCA sensor ranking algorithm.

Above result is for a single subject (i.e. subject 1). It shows that for all fingers, distal sensors are critical sensors. Unlike other studies (Kang et al., 2012; Chang et al., 2008; Wheatland et al., 2013) our method shows that placement of the sensor on the thumb is also critical. The sensor on the ring finger distal and middle phalange are equally important.

For cutting off the least important sensors based on PCA ranking algorithm we performed an analysis similar to the previous chapter. Sensors were placed in ascending order based on PCA ranking algorithm and the first derivative of *sensor importance* curve was computed. We observed the peak at 6^th^ place on the derivative curve as shown in figure 6 below.

**Figure 6:**
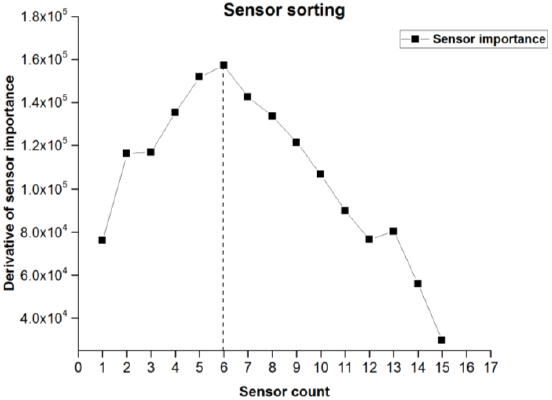
Sensor sorting. The derivative curve of sensor importance from PCA ranking algorithm. Selected sensors are shown below in figure 7(a) with a solid circle and missing sensors in figure 7(b) with a hollow circle. Thumb sensors were also selected unlike in previous studies (Wheatland et. al. 2013).

**Figure 7:**
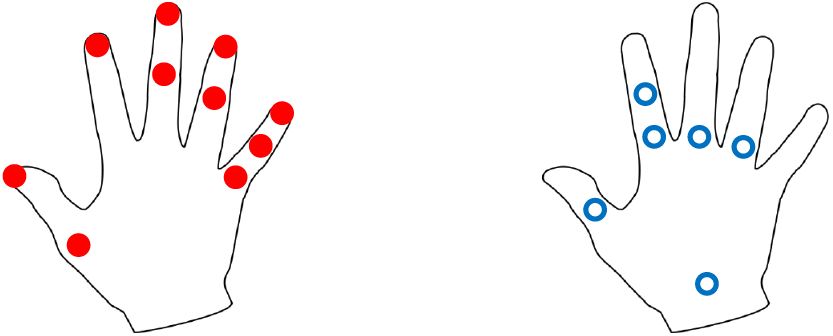
**(a)** position of selected sensors. All distal sensor locations were selected. Thumb sensors were also selected unlike in previous findings. **(b)** the position of missing sensors. Due to least variance in the sensor at the wrist, reference sensor was part of missing sensor group. All the sensors on proximal phalangeal digits were not selected due to less variance at proximal phalanx in comparison to another phalanx.

The peak value of sensor derivative curve was observed at 6^th^ place. The right-hand side of the peak, sensors were selected for computing all the joint angles. First 6 sensors were removed from the analysis as missing sensors.

The analysis was performed for all 12 subjects and ranking was generated for all sensors based on their PCA ranking score. Sensor contribution was computed for all sensors averaged across subjects. Results based on PCA ranking of orientation data across 12 subjects are presented in figure 8.

**Figure 8:**
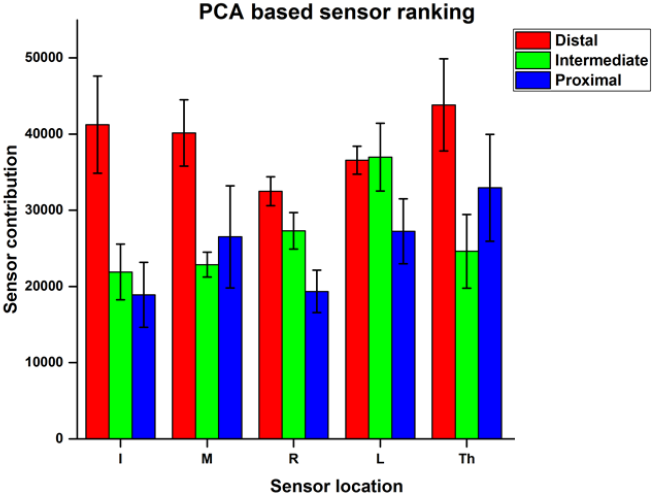
Sensor contribution across subjects. Bar graph represents the importance of each sensor across subjects. Error bar represents standard error of mean across subjects. The graph shows sensors located on distal phalanges are more important in comparison with other sensors.

From the above graph, we observe that, in general, distal sensors are relatively more important in comparison with sensors placed on other locations. However, the result varies across subjects. Contribution of sensors for each subject was computed separately. Peak value of derivative curve differs between subjects. Results for each subject, the derivative curve of contribution and selection of anatomical location are presented in table 1 below.

**Table 1:**
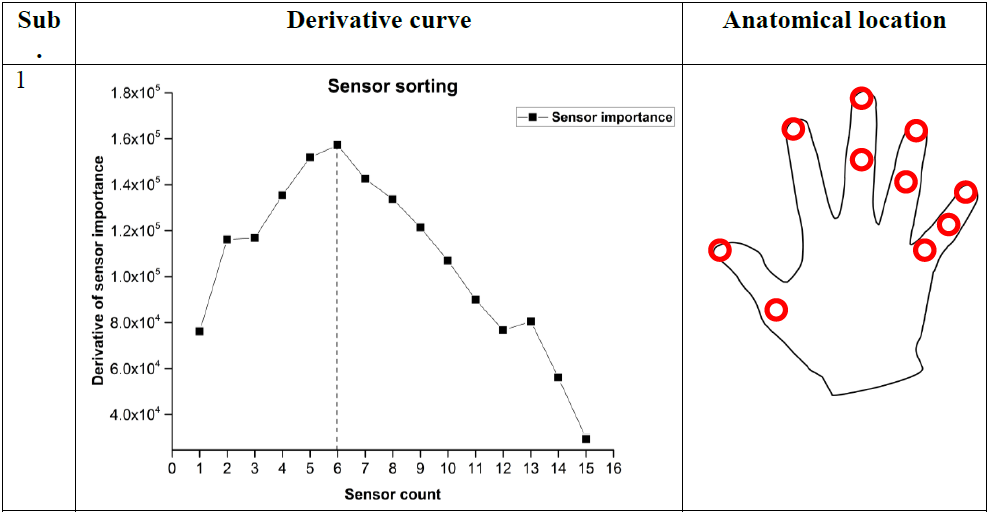
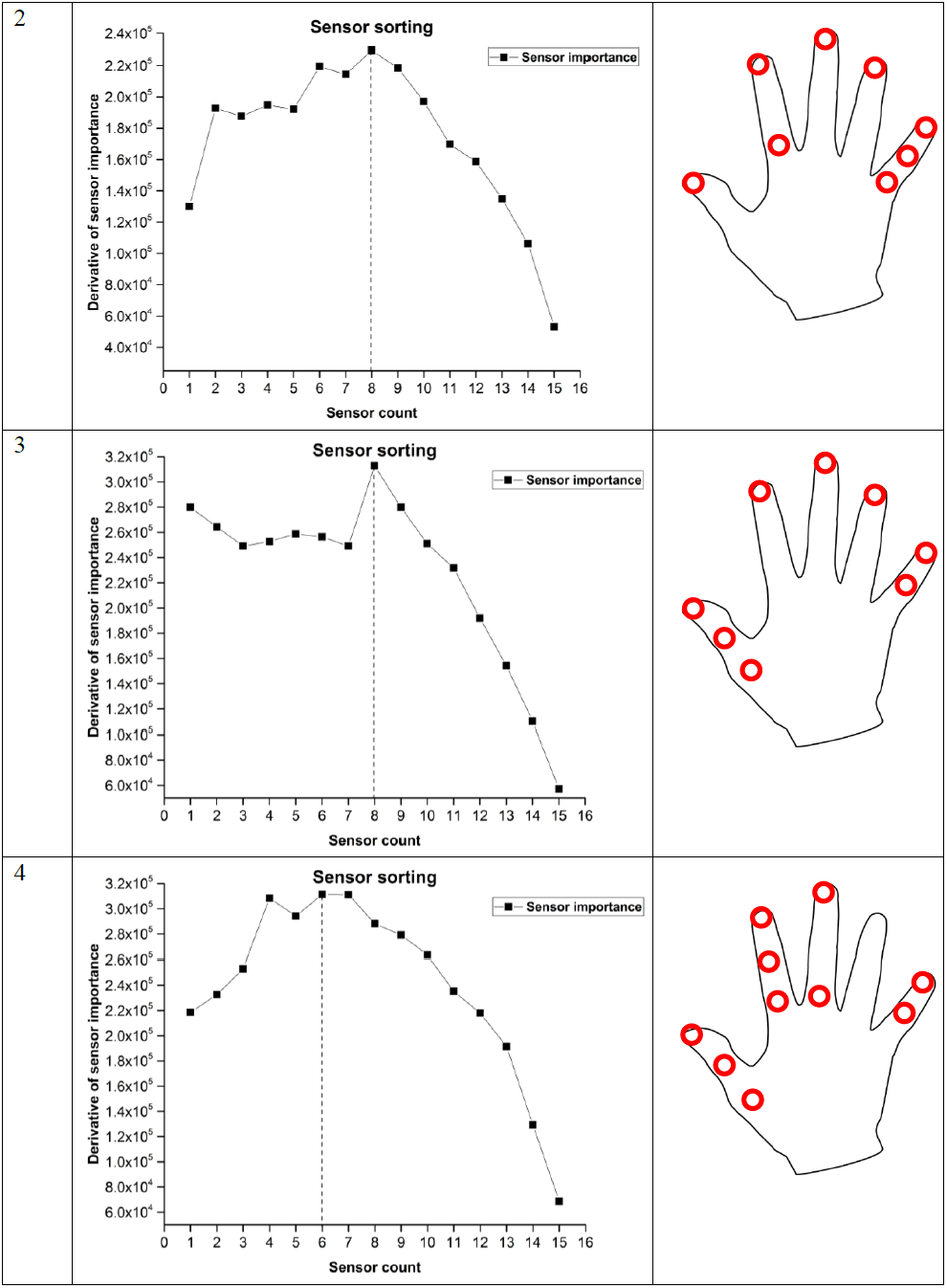
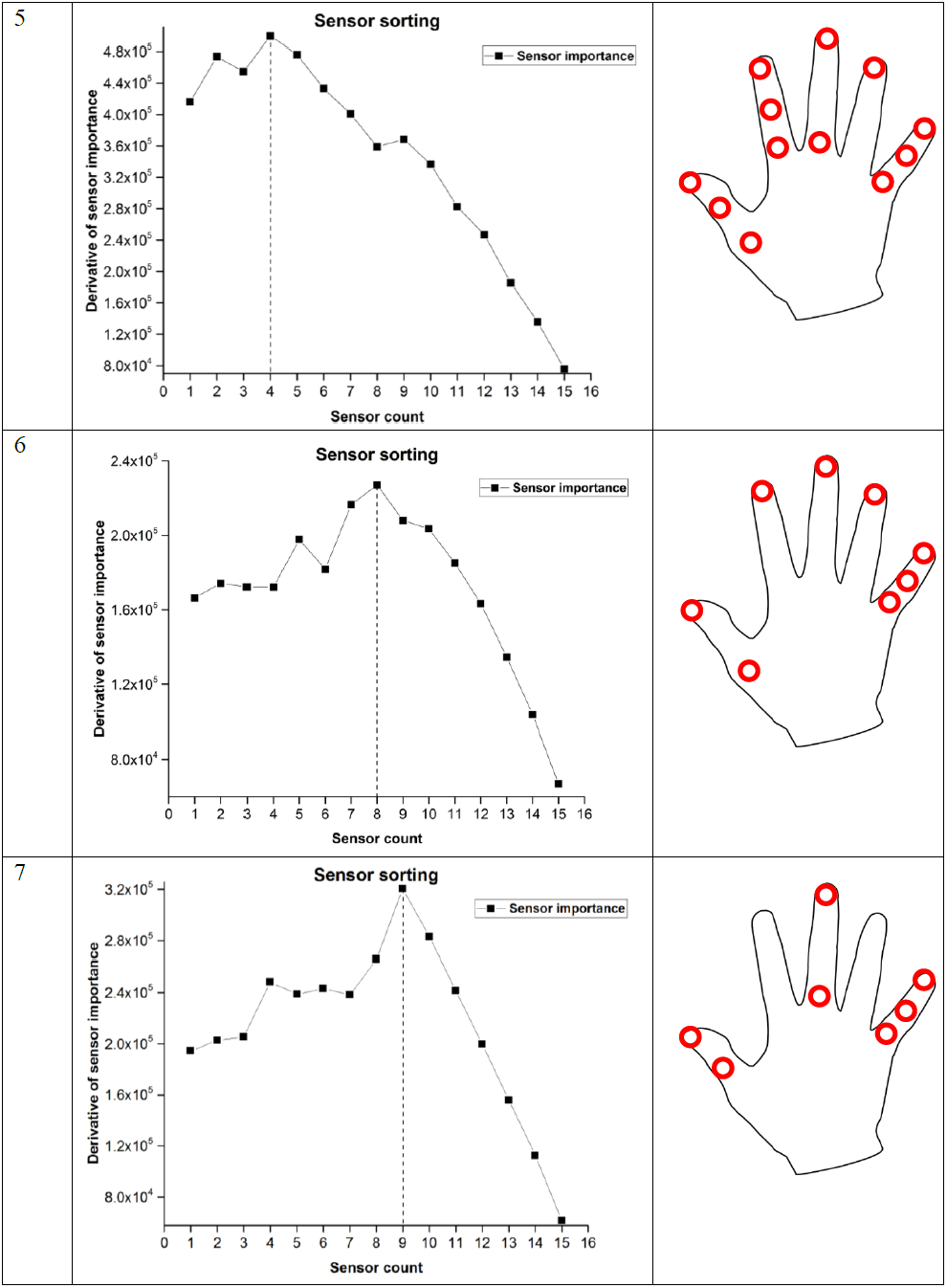
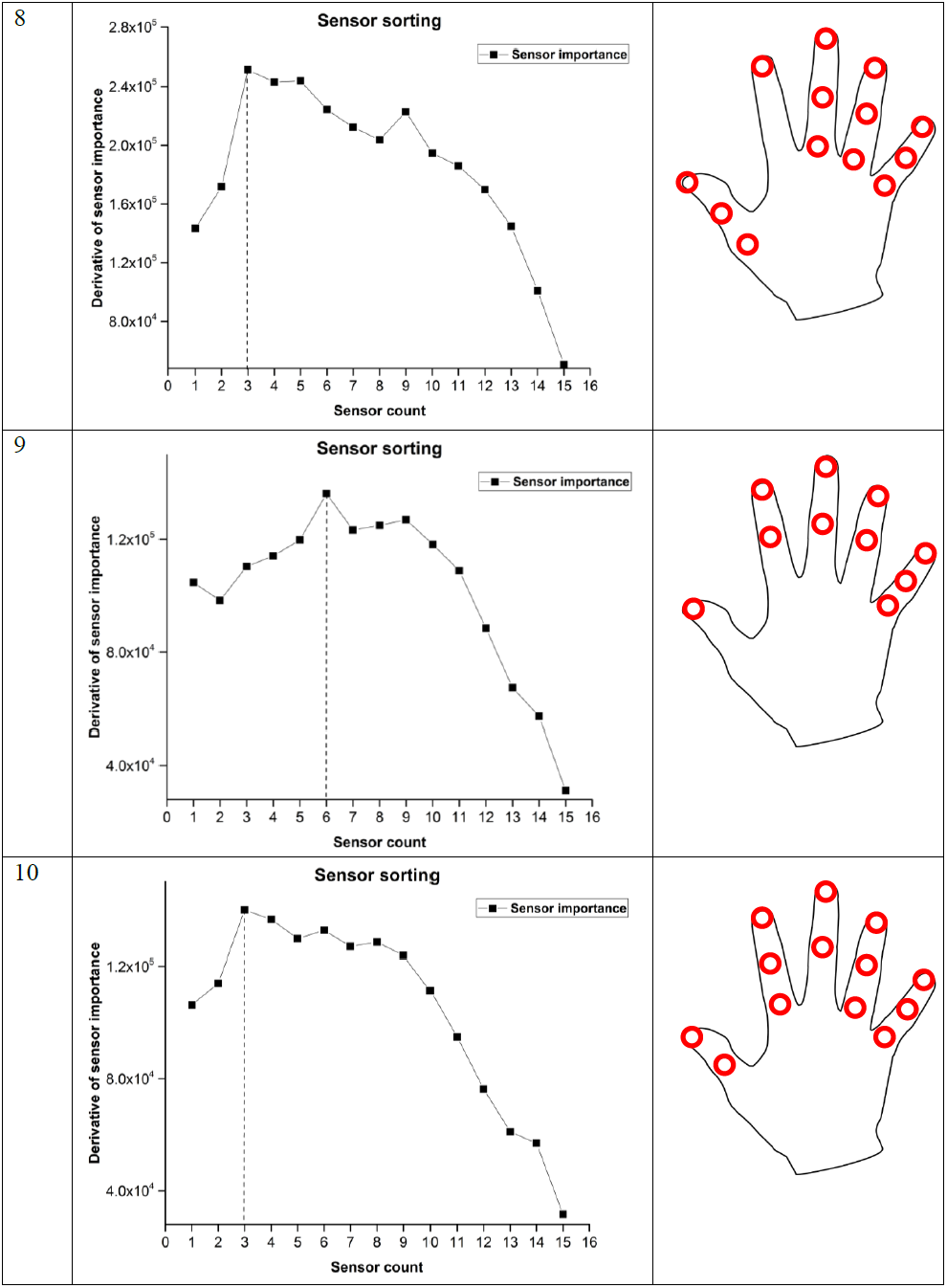
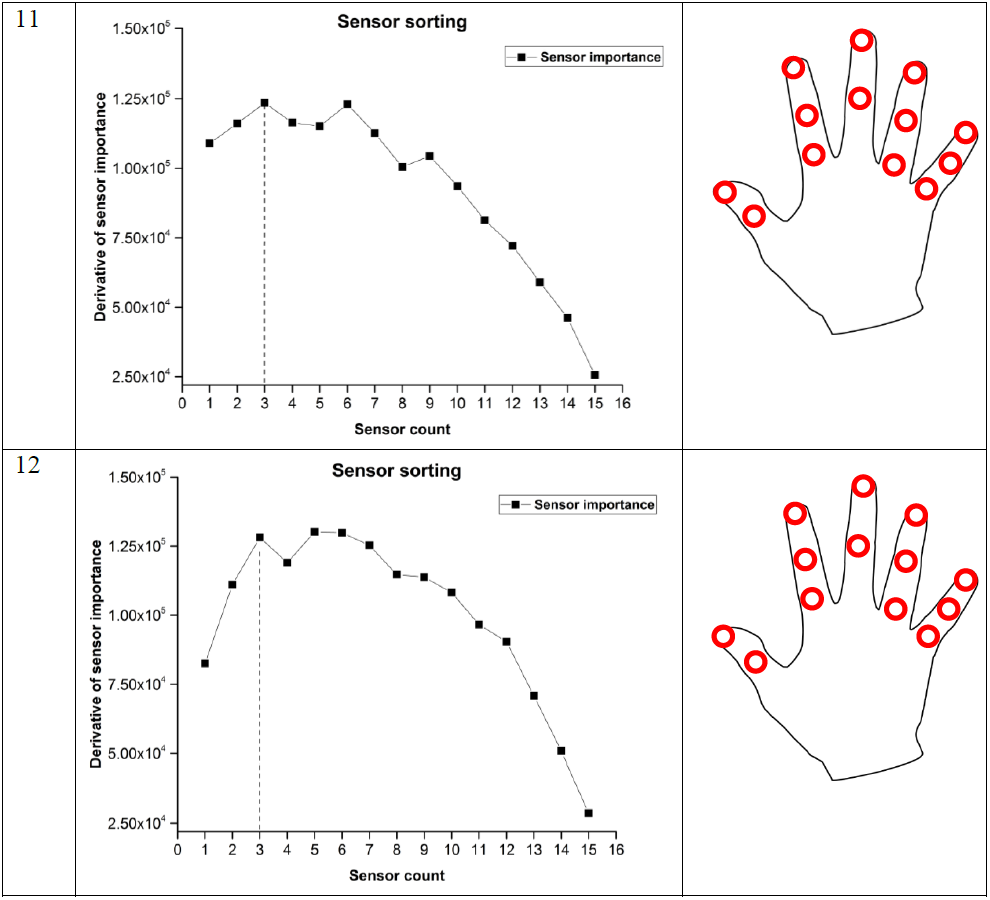
First derivative of sensor contribution curve and anatomical location of sensors placement.

This algorithm is for preliminary selection of the sensors from all the 16 sensors. For further reduction in sensor count detailed analysis is required. Our goal is to predict joint angles with reduced set of sensors.

### 1.2 Sensor correlation method

Correlation between sensors is an important feature for determining redundancy in the available set of sensors. Correlation between two signals can be easily found when it holds single signal value. Pearson’s correlation can be found which fits the straight line with slope value determining the goodness of fit and value of correlation coefficient lies between -1 and 1. If the correlation coefficient is positive, it shows two signals are positively correlated, if one signal increases another signal also increases. If the value of correlation coefficient is negative, then if one signal increases, other signal decreases. Zero value of correlation coefficient indicates no relation between given two signals. However, Pearson’s correlation coefficients cannot address nonlinearity in the signal. For addressing nonlinear characteristic of joint angles we used Spearman rank correlation coefficient which measures direction and strength of two signals between features. It helps in discovering the monotonic relationship between two variables. However, the monotonic relationship is not a strict assumption for Spearman rank correlation.

Electromagnetic tracking sensor gives three positions (X, Y, and Z) and three orientation values (Azimuth, elevation, and roll) as output. Joint angles can be derived using principles of mechanics (Zatsirosky 2000). Based on orientation data, we need to determine the correlation between two sensors. Current approach provides three different correlation coefficients. These three correlation coefficients are with respect to three orientations orthogonal to each other. Due to constraints of mechanics, algebraic operations on coefficients are not exactly legal. Ignoring one of the orientation may mislead the relationship between sensors. Hence we have a need to address this issue. Our method accounts for these limitations and finds correlation that also considers physical significance of sensors.

#### Method

The correlation coefficient between two sensors for all orientations was computed using spearman’s rank correlation. If the sample size is ‘n’ and the difference in rank between two variables is ‘d’ then Spearman rank correlation can be found using the following:

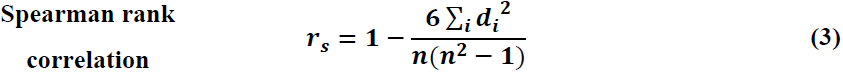

d_i_ is the difference between two ranks of features. n is a number of samples. A positive value of coefficient describes the nature of the relationship between two variables is positive. If the sign of correlation coefficient is negative, it indicates an inverse relationship between these two variables.

For computing correlation between all sixteen sensors, we used static postures with all 500 trials. Dataset consists of 35000 samples (70 samples per posture x 100 postures x 5 trials per posture). We computed Spearman rank correlation for each orientation separately. Hence we get three correlation matrices for a single subject corresponding to three different orientations (Azimuth, elevation, and roll). For determining the correlation between two sensors, we need to decide threshold value above which sensors we consider as “correlated” or “equivalent”. The threshold value of every orientation was computed using the following method. For each orientation, a matrix of correlation coefficient was converted into a vector and was arranged in ascending order (also called correlation curve). The first derivative of this *correlation curve* is computed and peak value was identified. Based on the value of peak corresponding value in correlation curve determines a threshold value for that particular orientation. The procedure is repeated for other two correlation matrices. Correlation curve across subjects and the first derivative of the curve are presented in figure 9(a) and 9(b).

**Figure 9:**
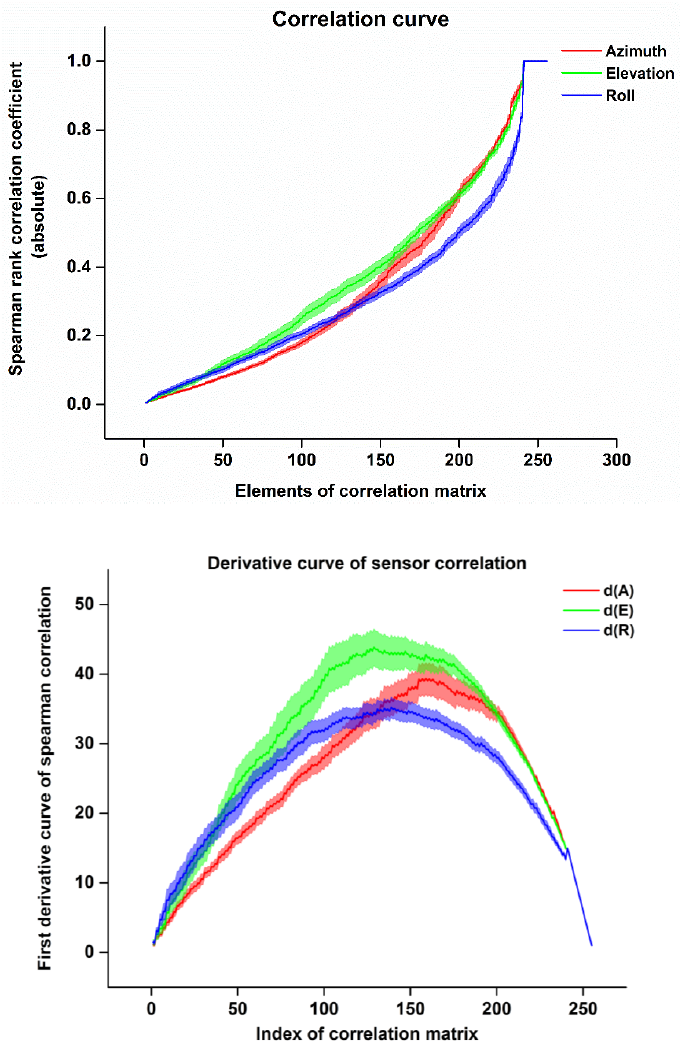
**(a)** Spearman correlation curve of the correlation matrix. Values of correlation coefficients are taken as absolute values. **(b)** The first derivative of correlation coefficient curve. The shaded area represents standard error of mean across subjects.

Figure 9(a) represents the absolute value of correlation coefficients of the correlation matrix for all orientations. We used the absolute value of correlation coefficients since we are interested in the magnitude of correlation rather than directionality. Figure 9(b) presents a first derivative curve of correlation curve. Peaks were observed at different places for different subjects. However, for all the subjects peak value for Azimuth is higher in comparison with elevation and roll. These peaks are used to identify threshold for determining the correlation between any two sensors.

For the purpose of illustration, single subject (i.e. subject 1) results are presented in figure 10.

**Figure 10:**
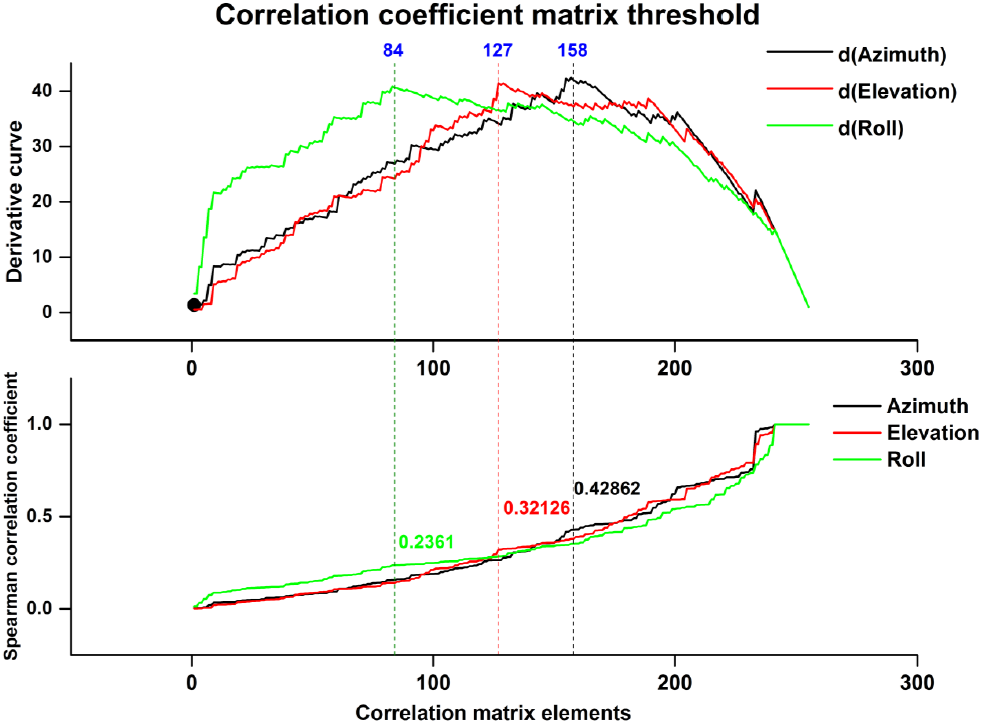
Correlation coefficient threshold points. **(a)** The derivative curve of sensor correlation curve for three different orientations. **(b)** Absolute of correlation curve were arranged in ascending order for all three different orientations. Peak values for the derivative curve were observed at different points (158, 127, 84) for (Azimuth, Elevation, Roll) respectively. Corresponding points are marked on absolute correlation curve.

The peak of the first derivative was computed and corresponding values were selected as a threshold on correlation coefficient curve. We observed highest value is corresponding to azimuth orientation which is mainly involved in flexion-extension movements. Hence these threshold values have physical interpretation from a kinematics prospective. Three different threshold values (0.4286, 0.3213, 0.2361) were calculated for this subject. Values above this threshold are considered to be a correlation between sensors. Three correlation matrices were computed corresponding to three orientations as shown in figure 11. From figure 11(a), we observe that few sensors are correlated with each other. However, for other orientations, many sensors are correlated. We found sensors that are correlated in all three directions shown in figure 11(d).

**Figure 11:**
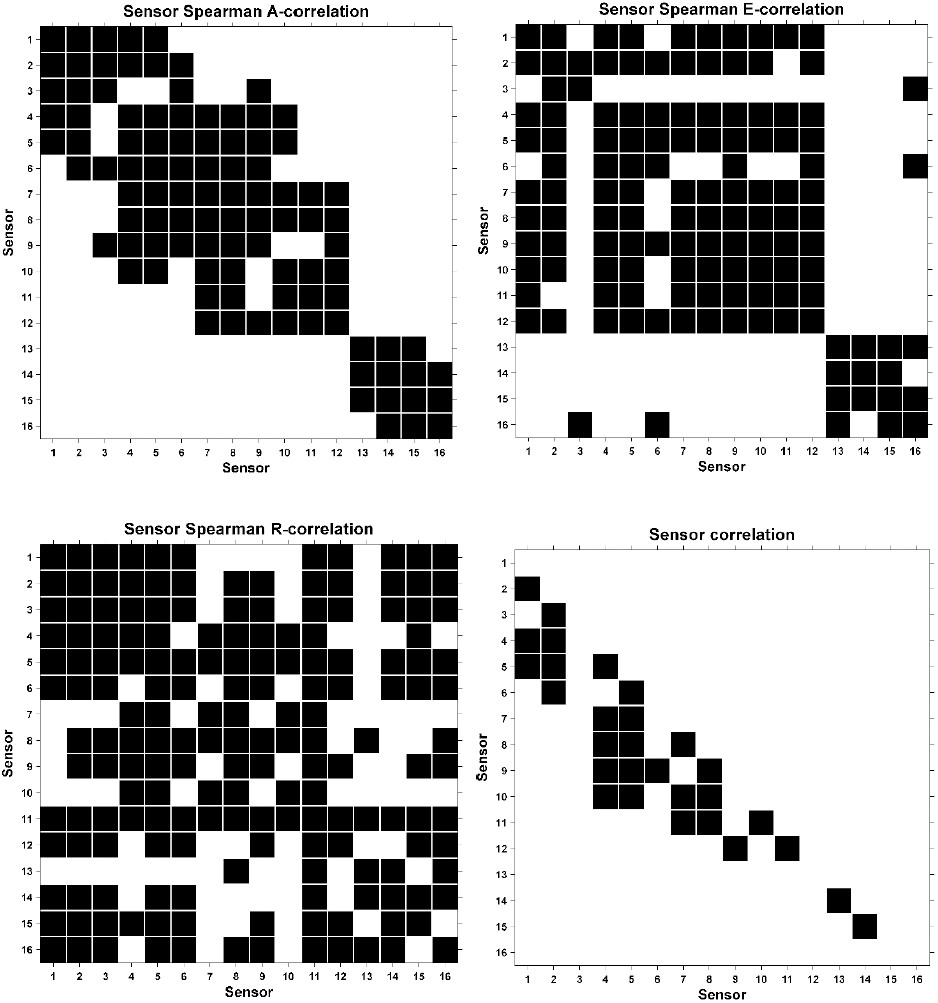
Sensor correlation matrices. Black color represents the correlation between two sensors. The size of the matrix is 16 × 16 gives correlation between all 16 sensors. **(a)** Correlation matrix between sensors in the azimuth direction. **(b)** Correlation matrix between sensors in the elevation direction. **(c)** Correlation matrix between sensors in the roll direction. **(d)** Based on the interaction of A-correlation matrix, E-correlation matrix, and R-correlation matrix, overall sensor correlation was computed.Correlation between sensors is shown in figure 11(d) is completely based on orientation dependent threshold values. We eliminated self-correlation of sensors (i.e. diagonal elements were removed from correlation matrix). The same analysis was repeated for all the subjects and correlation matrices were generated for all 12 subjects. Overall sensor correlation was observed and results are presented in **Appendix C**. For a single subject, the correlation between sensors is presented in table 2.

**Table 2:**
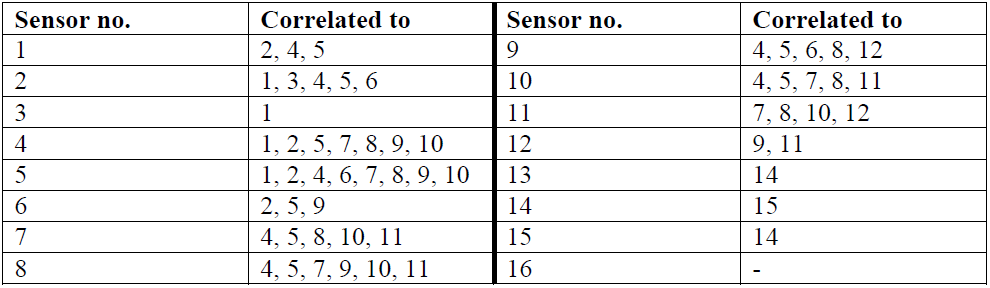
Sensor correlation for a single subject.

From above result, we note that sensors mounted on middle, ring and little fingers (i.e. 4, 5, 6, 7, 8, 9, 10, 11, 12) are correlated with each other. Correlation analysis was performed for all subjects and was normalized for 12 subjects. The result is plotted in figure 12 below.

**Figure 12:**
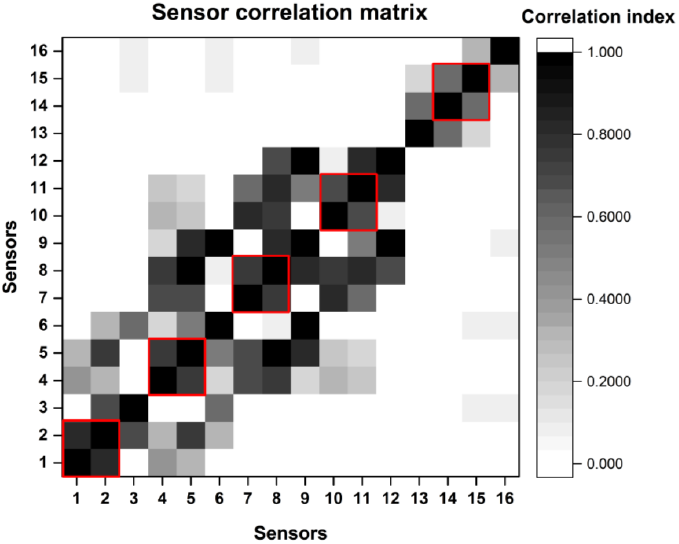
Sensor correlation matrix across subjects. Correlation between all 16 sensors was computed and normalized across subjects.

Figure 12 shows the correlation between all sixteen sensors normalized across subjects. Highlighted red color box represents the strong correlation between a distal and middle segment of every finger. This result is consistent across subjects. Our method presents a novel approach for determining the correlation between two sensors based on each orientation separately and shows consistency in results across all 12 subjects.

### 1.3 Decider network

Neural network can be a potential candidate for studying finger movements and hand function due to nonlinearity in the structure. We investigate the performance of a decider network, which is a neural network with single hidden layer and trained using Levenberg-Marquardt algorithm to minimize joint angle prediction error (i.e. network performance). These results are further used for quantification of sensor importance. If prediction error is large due to the presence of a particular sensor, that particular sensor is important. This error is chosen as a measure for sensor ranking.

#### 1.3.1 Network design

In the neural network, nodes in any layer take an input from the nodes in previous layer and Data from the previous layers are weighted based on strength of connection and summed before adding bias and passing through a non-linear activation function. Hence, data propagates from input layer (IL) to hidden layer (HL) to output layer (OL).

The output of the neural network is combinations of many nonlinear transformations of original input data (i.e. orientation data). Before finalizing network configuration, a set of parameters were empirically chosen.

*Number of hidden layers*: Number of hidden layers determines the amount of nonlinearity introduced for the dataset. If a number of hidden layers are more, the amount of nonlinearity on the dataset is large. However, large nonlinearity may suffer from overfitting.

*Number of nodes per layer*: This determines network architecture. The network could be made either converging or diverging or both. In general, networks are designed as convergent since inputs are expected to combine in a compact manner and transfer to next higher level abstraction.

#### 1.3.2 Network Training

Once the network is empirically designed, weights and bias values are initialized randomly. Dataset is mainly divided into two categories. First, internally constrained hand posture, Second, externally constrained hand postures. Internally constrained hand postures consists of hand postures without object manipulation (includes ASL and Bharatanatyam) a total of 70 postures. Externally constrained hand postures include object manipulation, a total of 30 postures. Two networks were trained to handle these two different datasets separately. The input to the network is orientation from the 16 sensors (no. of samples x dimensions = 168000 × 48) for internally constrained postures (5 trials per posture x 70 postures x 480 samples per posture). Similarly, for externally constrained hand postures with all 16 sensors size of input dataset is 72000 samples (5 trials per posture x 30 postures x 480 samples per posture) x 48 dimensions. The output of the network is joint angles. Since we have considered hand model to be 21 DoF, a number of nodes in output layer are 21.

The network cost function is mean squared error between actual and predicted joint angles. The function that needs to be minimized is the following form:

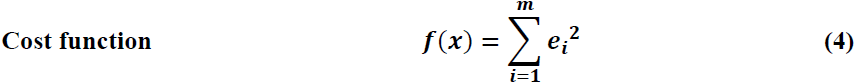

Function f is cost function which needs to be minimized. e represents residual between predicted and actual joint angles. *m* represents number of samples in the dataset.

Our objective is to minimize the error between predicted and actual joint angles. We chose Levenberg-Marquardt (LM) algorithm for updating weights and bias values since it is a fast converging algorithm. It combines the advantage of Gauss-newton and gradient descent algorithm for faster convergence based on the properties of error surface. In general, Newton method is used as an alternative method for conjugate gradient methods for fast optimization. This method uses Hessian matrix for updating weights. However, computing Hessian matrix and its inverse are computationally expensive. Hence, instead of calculating Hessian matrix directly, it can be derived from Jacobian matrix using following equation

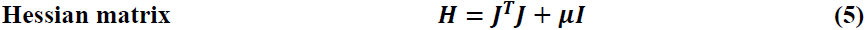

The Jacobian matrix can be computed as derivative of errors w.r.t parameters (weights), μ represents damping factor and *I* represents identity matrix.

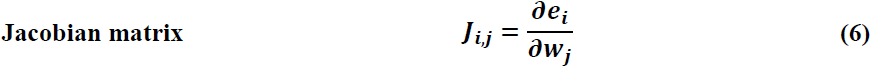

Where i, j represents number of training samples and number of weights in the network respectively. The size of the Jacobian matrix increases with increase in number of samples. However, Jacobian is less complex than Hessian matrix. The gradient vector is computed as follows:

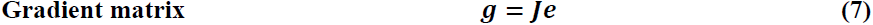

e is a vector of errors presented in the network. The weights are updated based on the following equation

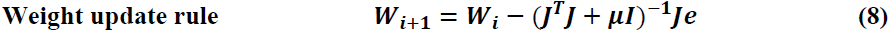

If μ is zero, weight update rule is same as Newton method using approximate Hessian matrix. If μ is large enough, the method becomes gradient descent with smaller learning rate. Newton method is more accurate when the loss is lesser or near convergence. The value of damping factor is initialized to be large so it works as gradient descent algorithm and as the value of loss function reduces, μ decreases.

##### LM algorithm and weight update rule step by step

*Step1:* Initialization of algorithm, weights are initialized randomly. Compute the error E, as the difference between actual and predicted values.

*Step 2:* Compute the Jacobian matrix using equation (6) as a change of error with respect to weight.

*Step 3:* update the weights based on equation (8) and evaluate total error.

*Step 4:* If the current error is increased as a result of weight update, reset back the weights to the previous value and increase the value of μ by some factor and jump to step 3 again.

*Step 5:* If current error is lower due to weight update, then accept the current weights, reduce the value of μ and jump to step 3.

*Step 6:* Repeat the steps until the actual error is lesser than desired value.

One major drawback of this algorithm is the size of a Jacobian. If the number of samples are many, then it is computationally intensive to compute Jacobian. However, Jacobian can be divided into two equal matrices as follows:

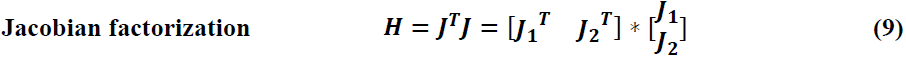

From the above equation, it is not necessary to store all the samples at a time. The Hessian approximation can be computed by summing all individual subterm Jacobians. The method of reducing computational load is helpful for reducing computational power.

#### 1.3.3 Joint angle prediction

The input to the network was orientation data (168000 samples x 48 dimensions). The output of the network was 21 joint angles. The network was optimized using Levenberg-Marquardt optimization backpropagation algorithm. Number of nodes in hidden layers was optimized empirically for three subjects. Thirty nodes in hidden layer were able to generate satisfactory performance. Neural network toolbox was used for designing and for configuration purpose in MATLAB. We performed analysis for all the twelve subjects. Network performance was measured based on *Mean squared error (i.e. MSE)* in prediction.

Mean squared error was calculated for test dataset for all 12 subjects. The input to the network was 16 sensors orientation (48-dimensional dataset). For the purpose of physical interpretation, root mean squared error (RMSE) for test dataset across 12 subjects for all 21 joint angles are presented in figure 13

**Figure 13:**
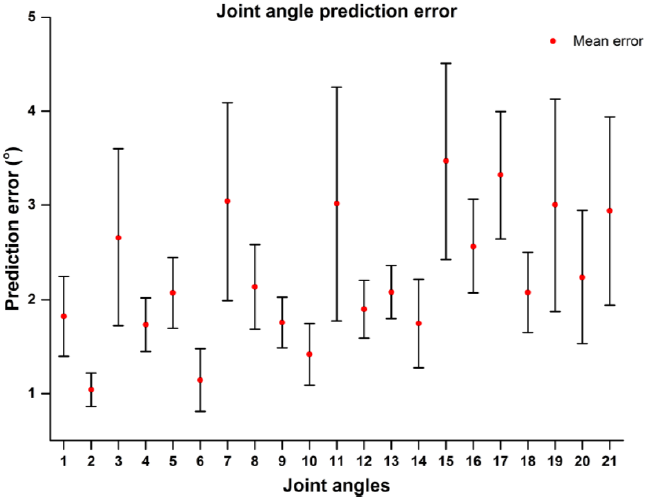
Joint angle prediction error (i.e. root mean squared error) for test dataset. Joint angle prediction shows large error in case of all 16 sensors presented for the single hidden layer with 30 nodes. Error bar represents standard error of mean across 12 subjects.

From the results presented in figure 13 and 14, we observe that for sensor ranking empirically chosen parameters for the network configuration performs better in terms of joint angle prediction. We selected the performance of this network for further ranking the sensors.

**Figure 14:**
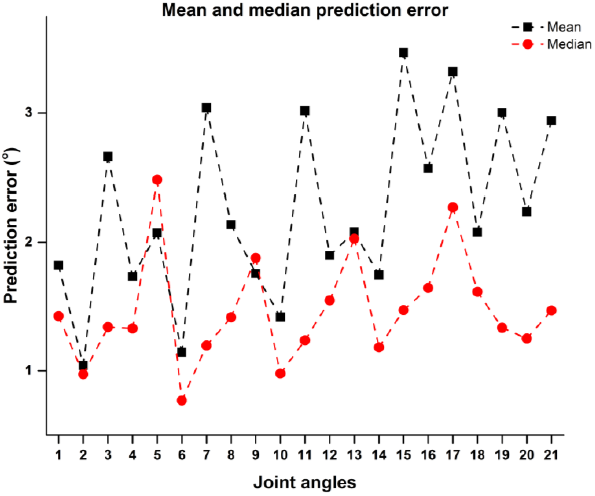
The mean and median of joint angle prediction error. Mean value of predicted joint angles is larger than the median value of predicted joint angles. Larger mean values are possibly due to large prediction error in some subjects. For other subjects prediction error lesser than 1.5° result in lower median values of joint angle prediction.

#### 1.3.4 Sensor ranking based on network performance

For ranking the sensors based on the performance, we modified the *backward selection* algorithm. For finding the importance of a sensor, we made network input to be zero for a particular sensor and tested network again on the training dataset. The increase in the network performance error is an indirect measure of sensor contribution to overall postures. If the MSE is higher by making values zero for a particular sensor, it means that the sensor is contributing relatively more in comparison with other sensors. For every subject, we performed sensor backward-selection separately. For illustration purpose, single subject (i.e. subject 1) result is presented in figure 15. For rest of the subjects, results are presented in **Appendix D**.

**Figure 15:**
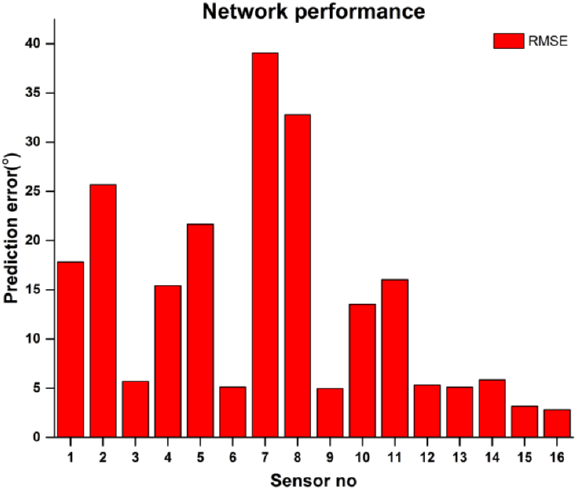
Sensor-based performance measure (subject specific) (i.e. subject 1). Prediction error due to the absence of the particular sensor. A sensor placed on ring finger distal and middle phalanges show large prediction errors.

Sensors were ranked based on the performance measure for every subject separately. Network performance was tested for training dataset after removing a particular sensor.

### 1.4 Sensor selection based on systematic approach using PCA ranking, sensor correlation, and network performance

Our method uses PCA ranking, sensor correlation and network performance for selection, removal or swapping of any sensor. Our method uses PCA ranking as preliminary sensor selection criteria. We choose sensor variance as primary criteria for selecting a sensor. Our approach for finding sensor correlation is second criteria used to determine redundant sensors. We measured neural network performance after training for finding corresponding importance for sensors. Large mean squared error corresponds to the higher importance of a sensor. Our third criteria was sensor performance in decider network.

#### Method

First, PCA ranking was performed on all the sensors and the first derivative of sensor importance curve was computed. The peak of the first derivative curve is used for preliminary sensor selection. Sensors were divided into two groups 1. Selected sensors 2. Unselected sensors.

Further, we explored the possibility of unselected sensors being correlated with selected sensors. For determining correlation we have used our novel approach for finding a correlation between two sensors. However, a particular unselected sensor might be correlated with sensors that are selected and also with sensors that are not selected. To resolve this ambiguity we compared the performance of the sensor in the neural network. If total mean squared error performance is larger for selected sensors in comparison with unselected sensors, the sensor is classified as correlated. However, if unselected sensor has more correlation with sensors that have a large mean squared error then we conclude that sensor is classified as uncorrelated. Unselected-correlated sensors were dropped out and we further proceed with unselected-uncorrelated sensors.

Highest performance error (using the neural network) for unselected-uncorrelated sensors were computed and compared with lowest performance error in selected sensors. If the largest error in uncorrelated-unselected sensors is smaller than the smallest performance error in selected sensors, unselected-uncorrelated sensors are dropped out. If performance error for the unselected-uncorrelated sensor is larger than smallest performance error in selected sensors, sensors were swapped. The procedure is repeated until all the uncorrelated sensors either become correlated with selected sensors or performance measure is smaller than the selected sensors.

We used a hard threshold bound for selecting number of sensors which is ‘8’. If the number of selected sensors are 8, our method is considered to have converged and presents selected sensors as best possible set of sensors. If number of sensors are greater than 8, we further use reduction algorithm for determining lowest important sensor and remove them. According to reduction algorithm, sensors with least performance error are selected and corresponding correlated sensors are selected. Total performance mean squared is computed. Sensors with least score are dropped till hard bound is reached (i.e. 8 sensors). In this case, we have selected threshold on a number of sensors to be ‘8’.

Block diagram of our sensor selection method is described in figure 16. Procedure for selecting the sensors was performed for each subject separately. Location of selected ‘8’ sensors for different subjects is shown in table 3.

**Table 3:**
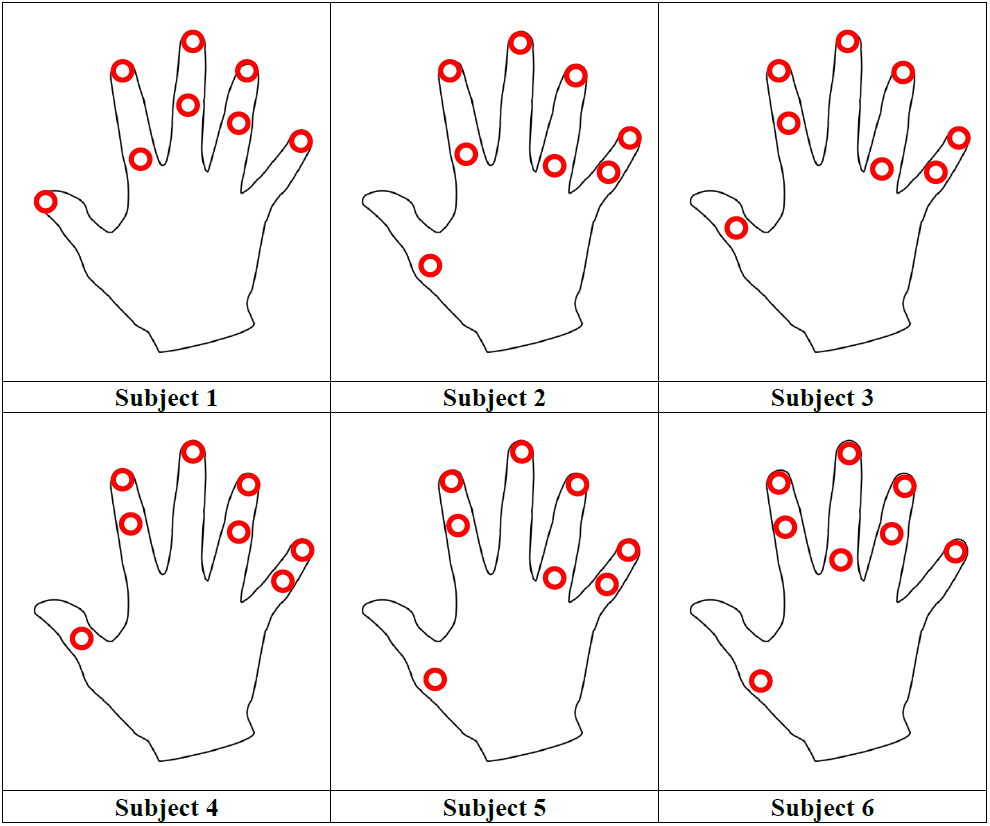
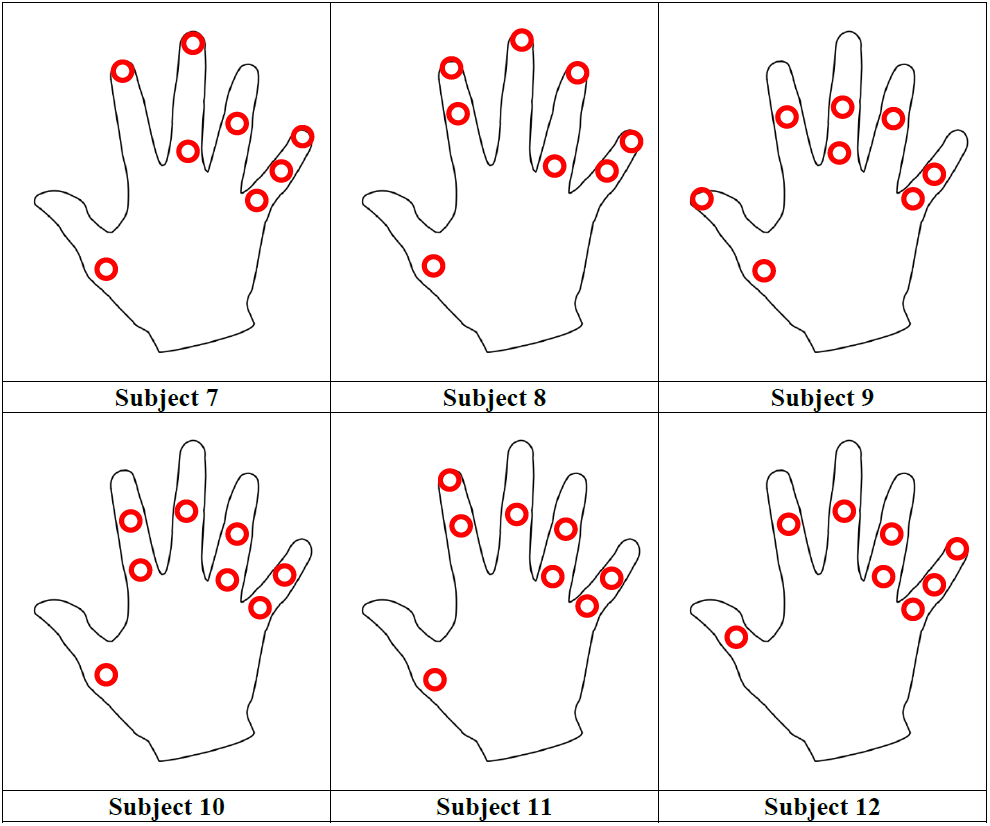
Subject specific sensor location.

**Figure 16:**
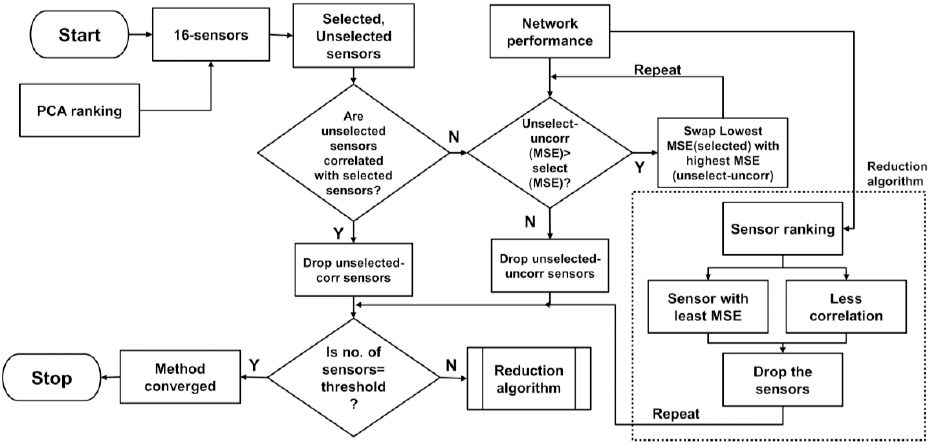
Sensor selection method. Block diagram shows the procedure for selecting a number of sensors. If a number of selected sensors are greater than ‘8’ (threshold), reduction algorithm needs to be applied on selected sensors for finding and removing redundant sensors.

### 1.5 Joint angle prediction using reduced set of sensors

Accurate joint angle prediction from reduced sensors is important from biomechanics point of view. Various approaches have been taken for joint angle reconstruction or joint angle prediction. Approach for predicting joint angles includes a method such as a neural network (Gorce et al., 2008; Mora et al., 2012) and data-driven approach for classification of various grasping patterns using random forest (Chang et al., 2007). In this thesis, we have used two basic data-driven approaches for accurate prediction of joint angles. We focused on *locally weighted regression* nonparametric method from selected sensors based on the output of PCA ranking algorithm and *neural network* as a parametric approach for joint angle prediction using our novel methodology. Topics are described individually in further subsections.

For joint angle prediction, our dataset is divided into two categories 1. Internally constrained (By intention) hand postures 2. Externally constrained (By objects). 1^st^ group consists of 168000 samples (70 postures x 5 trials per posture x 480 samples per trial) whereas 2^nd^ group consists of 72000 samples (30 objects x 5 trials per posture x 480 samples per trial) for a single subject data. In both cases, the dataset was divided into a training set (80%, internally constrained: 134000 samples; externally constrained: 57600 samples) and test set (20%, internally constrained: 33600; externally constrained: 14400 samples).

#### 1.5.1 Non-parametric approach

##### 1.5.1.1 Locally-weighted regression

Non-parametric models preserve the available training data and use every time for query point. One such memory based model can be used for joint angle prediction. This model is used as a regression model for prediction of joint angle query points based on the nearest (i.e. local) points. In the case of the nearest neighbor, equal weights are given and a number of neighbors are selected globally. To overcome this issue, a kernel function is used where weights are given based on distance. If neighbors are closer, they are given a higher score. The weighted function, is also called as kernel function. It compares and gives similarity between two examples. The simplest form of kernel is Gaussian kernel which is expressed in below in equation 10

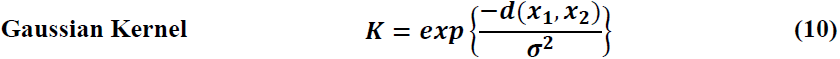

Where d(x1, x2) is the distance between two points x1 and x2. For Gaussian kernel, the distance between a query point and training sample point was computed using l2-norm. Gaussian kernel function was selected as a function for allocating weights. The width of kernel σ is an important parameter which selects how quickly weights fall off based on distance. If the distance between a query point and training sample point is small, weights will be closer to 1. If the distance between training point and query point is large, weights will be closer to 0.

Value of joint angles is predicted based on the following equation

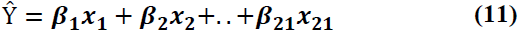

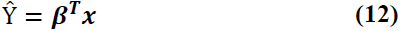

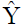 is predicted vector of joint angles. The objective of the cost function used in LWR (locally weighted regression) is to minimize the error between a predicted value (ŷ) and the output value (y) described below.

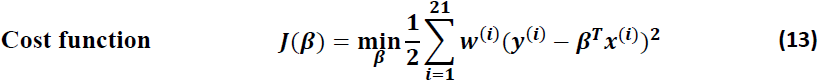

Where W is diagonal weight matrix. i varies from 1 to 21 for predicting hand joint angles. To find the minimum value of *β* that minimize the cost function, we differentiate equation (13) with respect to *β*. Closed-form solution is given as below

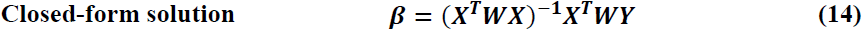

Joint angles are computed from reduced number of sensors. Locally weighted regression is used for computing all 21 joint angles of hand from a reduced set of sensors based on the distance of a query point from the points available in input space.

We optimized the kernel width parameter based on 5-fold cross validation for 3 different subjects and found that 0.5 is optimum kernel width. For rest of the subjects, we selected value of kernel function as 0.5, for joint angle prediction.

Dynamic postures were considered for generating a large pool of data points. Dynamic postures are generated by dropping out first 500ms data (60 points) and last 500ms data (60 points). Total 480 points per posture were considered. The model was tested on two datasets, 1. Externally constrained and 2. Internally constrained separately and performance is reported. Dimensionality of input data to LWR model is 3N, where N varies based on a number of selected sensors. The value of N varies depending on PCA sensor ranking algorithm. For different subject selected number of sensors were different. Results are presented for all the subjects separately in table 4.

**Table 4:**
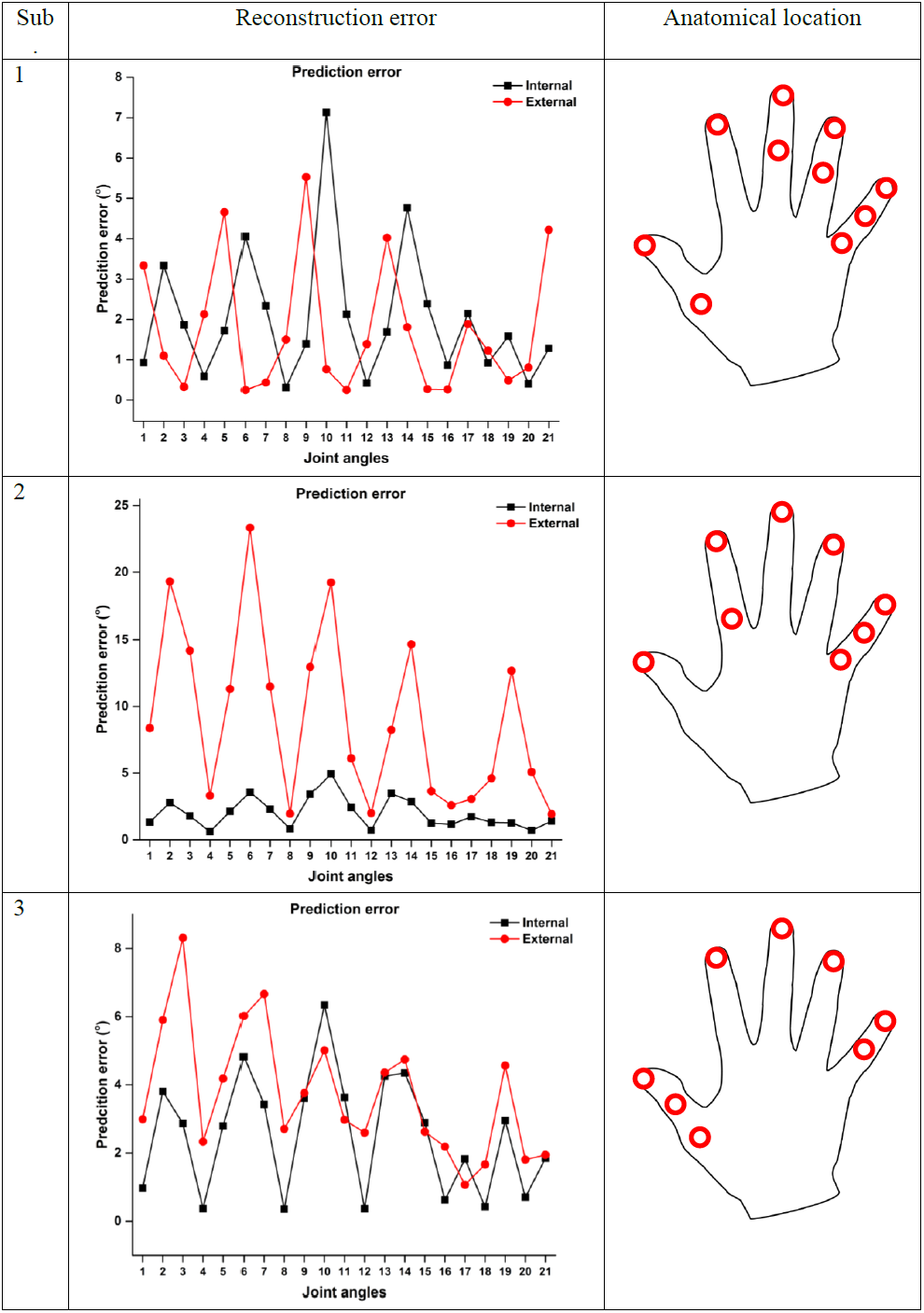
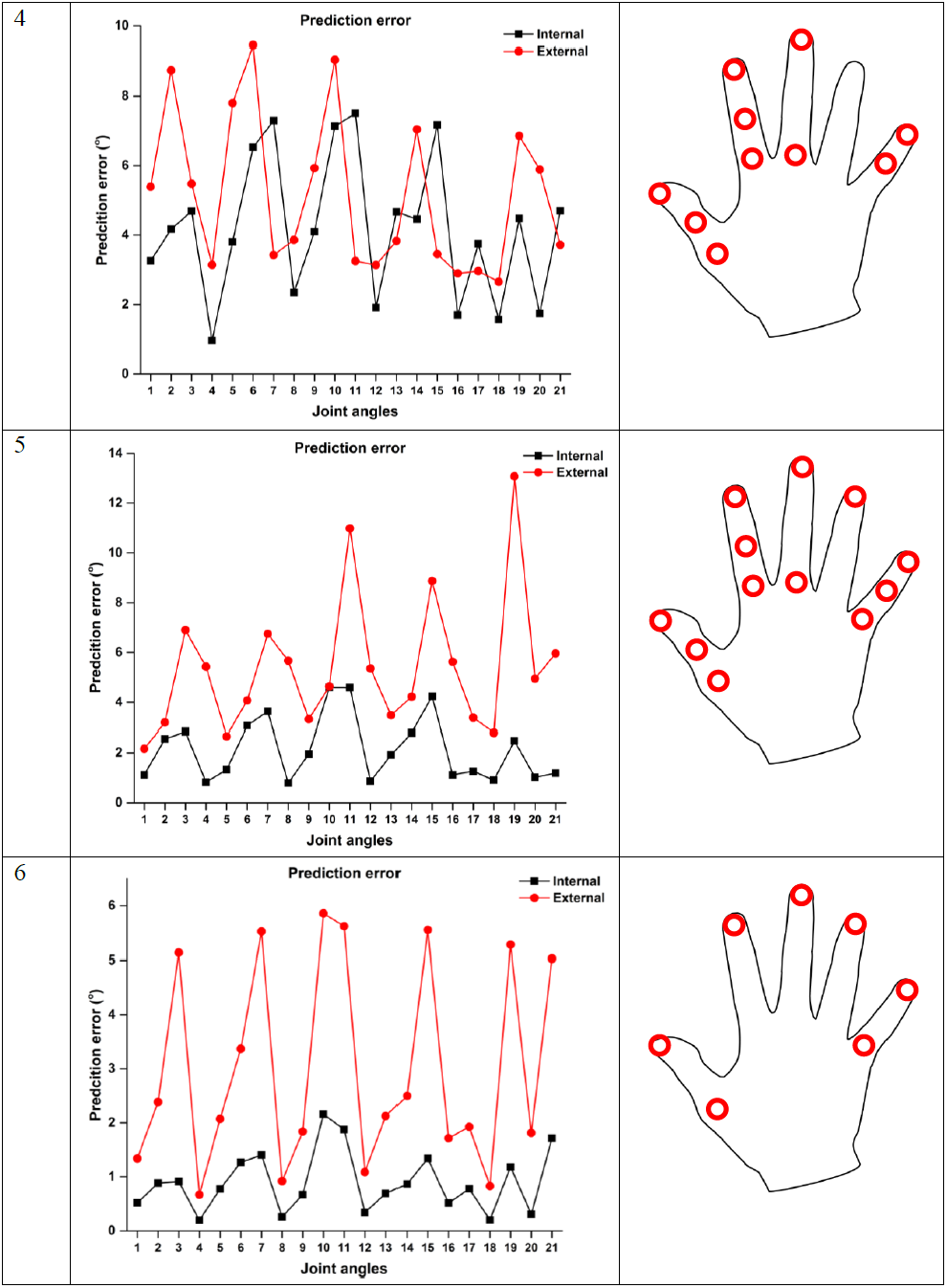
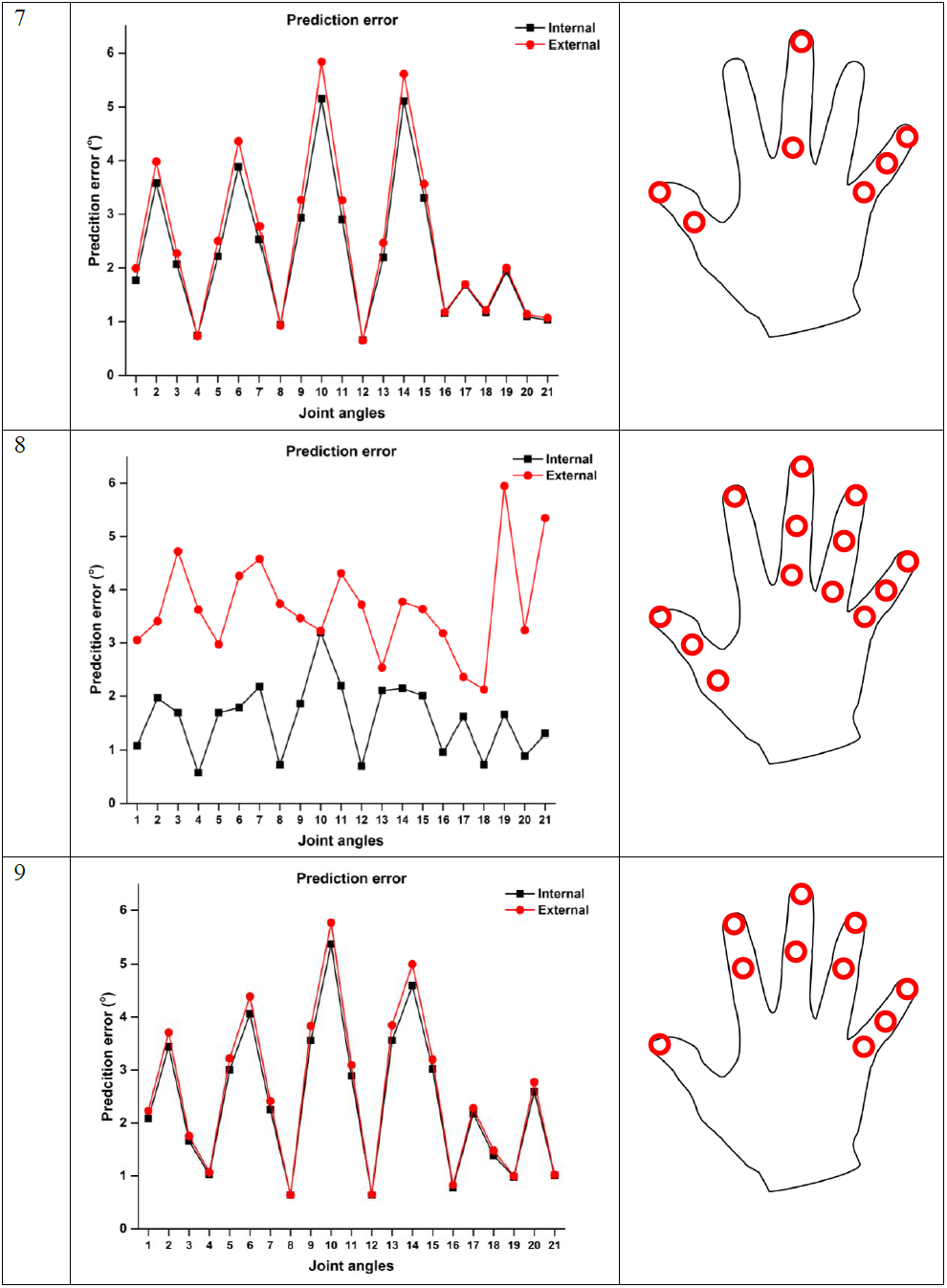
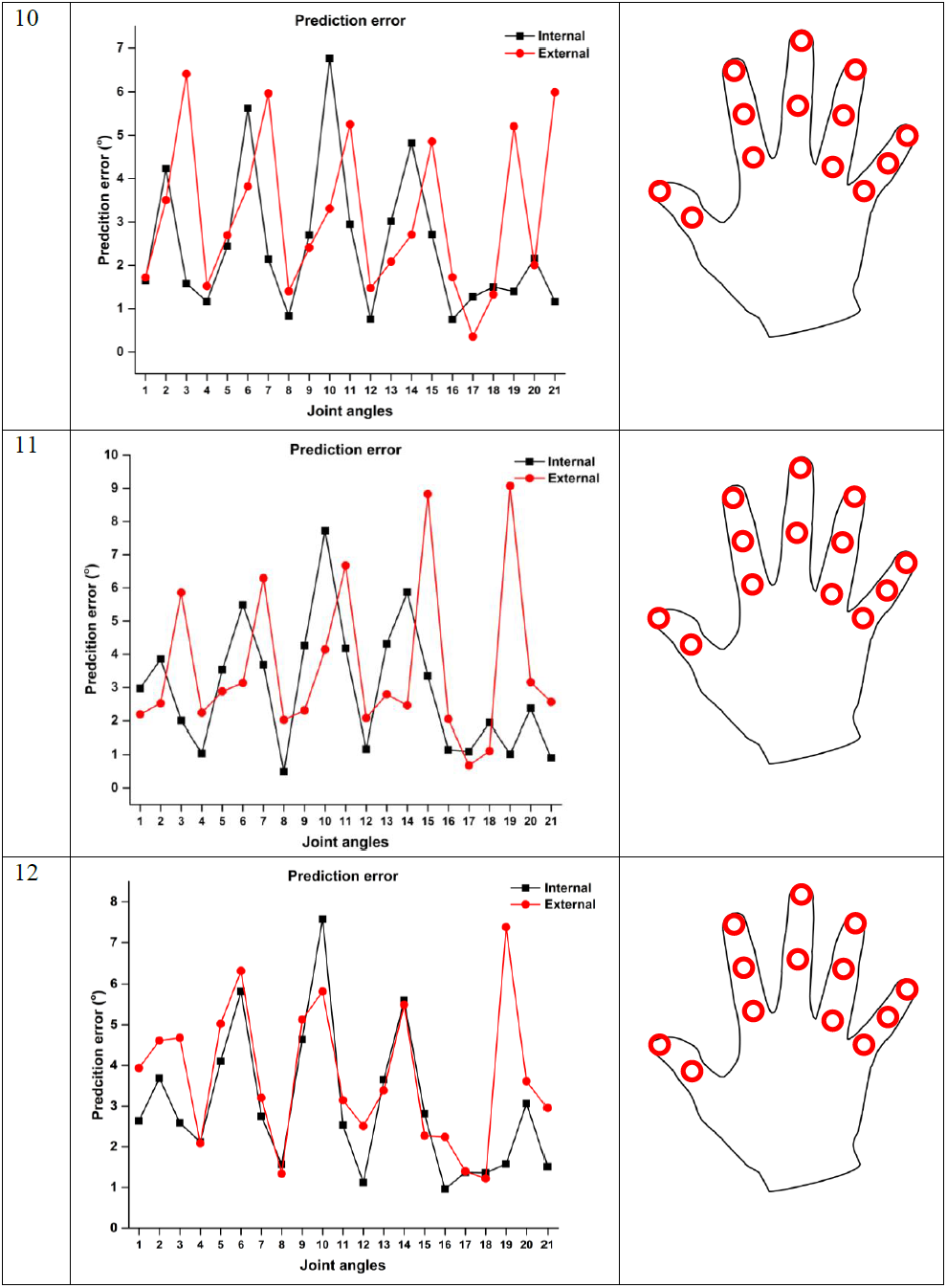
Joint angle reconstruction error with a reduced set of sensors for the internally constrained postures and externally constrained postures.

Joint angles were reconstructed using lesser number of sensors ranked based on PCA ranking algorithm. For different subjects, number of sensors selected were different. Joint angle reconstruction errors are different. However, common observation shows that PIP joint angles of middle, ring and little fingers are showing higher joint angle error. Above table shows mean value of joint angles reconstruction error across 5-fold cross validation. LWR model was tested based on the selected sensors for each subject separately. Test error is computed and plotted across all 12 subjects as shown in figure 17 below.

**Figure 17:**
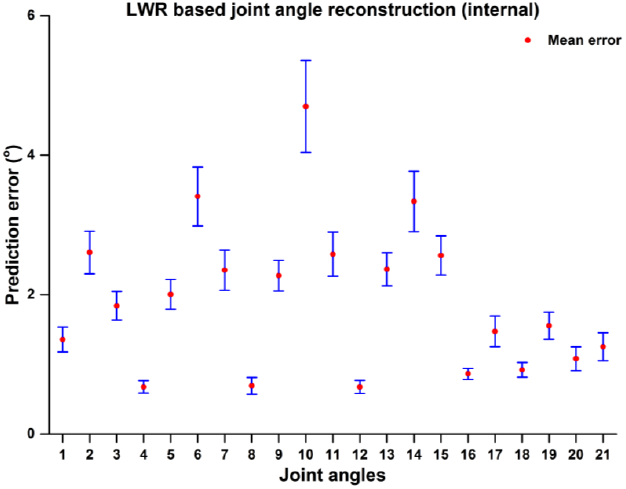
Joint angle reconstruction for internally constrained postures across all 12 subjects. Error bar represents standard error of mean across subjects.

Joint angles were reconstructed across subjects using LWR method and following result was found for test dataset.

**Figure 18:**
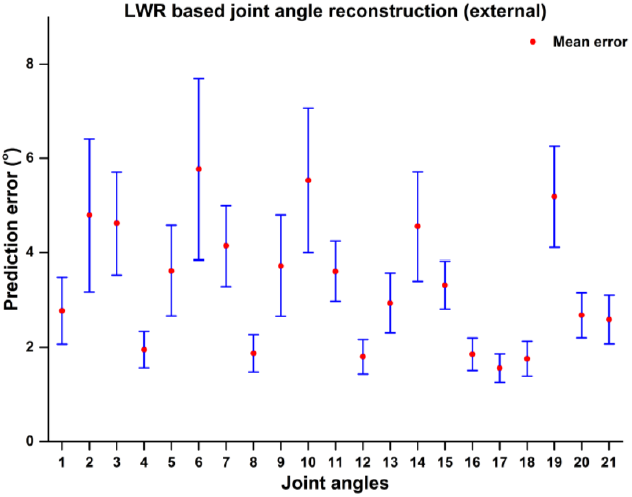
Joint angle reconstruction for externally constrained dataset using LWR across subjects. Error bar represents standard error of mean across subjects.

For externally constrained postures, joint angle reconstruction error for test dataset is relatively large, in comparison with internally constrained postures. Large variability is observed in PIP joint angle of the index, middle and ring finger.

**Figure 19:**
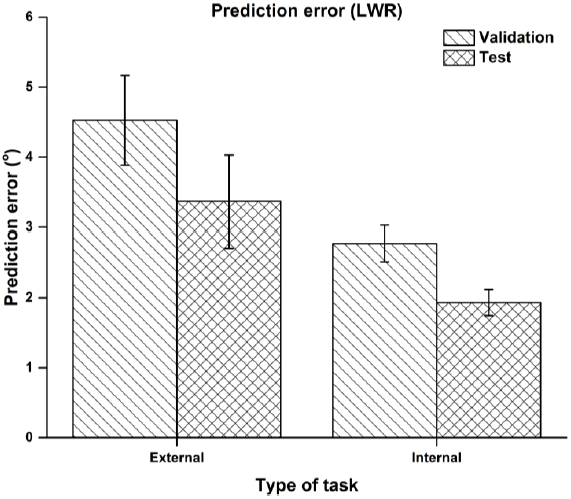
Joint angle reconstruction error across subject across joint angles (i.e. root mean squared error). Error bar represents standard error of mean across subjects.

### 1.6 Parametric approach

#### 1.6.1 Neural network model

We used neural network model for predicting joint angles from a reduced set of sensors. The network architecture consists of three layers, input (IL), hidden (HL) and an output layer (OL). The input layer has data from 8 selected sensors. Total number of input nodes are 24 (i.e. 8 sensors x 3 orientations) and output nodes are 21, fixed a priori. Other network parameters were optimized empirically. Following this, network weights are initialized randomly. Since joint angles are above the range of sigmoid and tangential neurons, linear neurons are used in the output layer. Training is done for minimizing prediction error between actual joint angle and predicted joint angle.

In one of the network configurations, number of hidden layers were chosen as ‘1’. There were 30 nodes present in the hidden layer. Levenberg-Marquardt algorithm was used for the purpose of updating weights of the network. Data from selected sensors were used for the purpose of training, and network performance was measured as mean squared error. However, for the physical interpretation of an error, the square root of the mean squared error was computed. The results are presented in table 5.

**Table 5:**
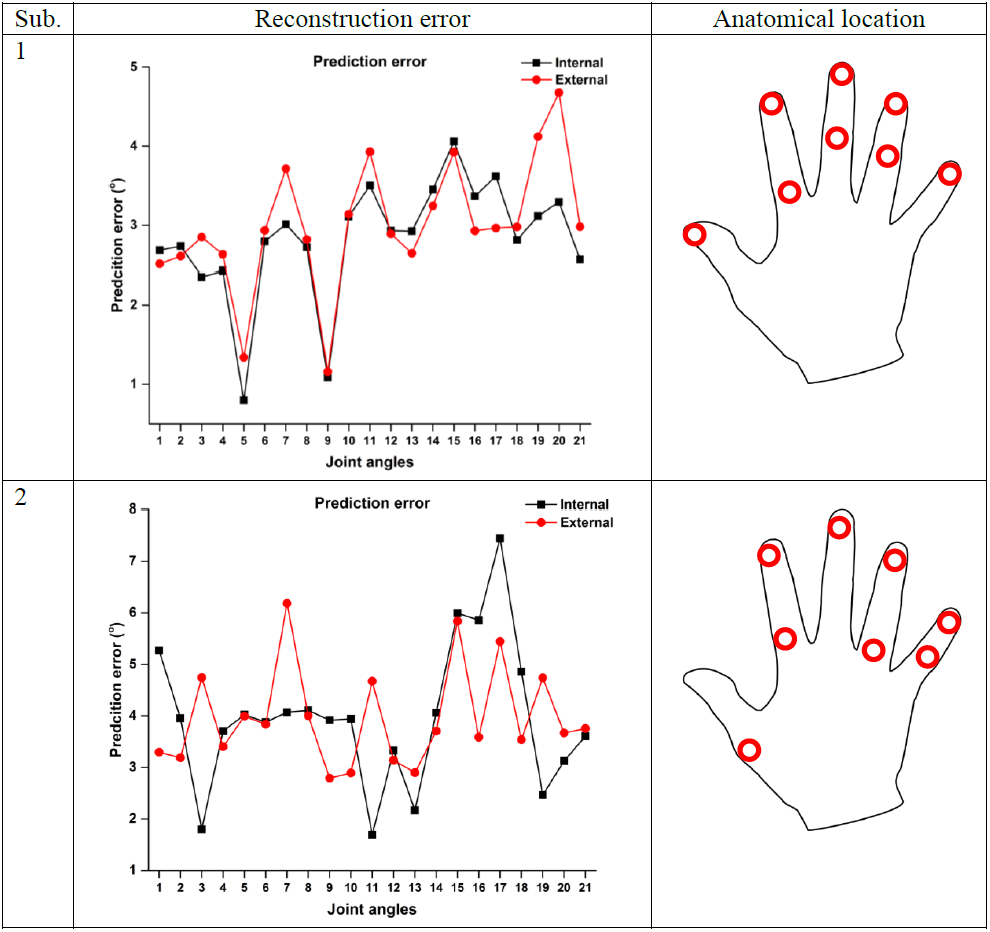
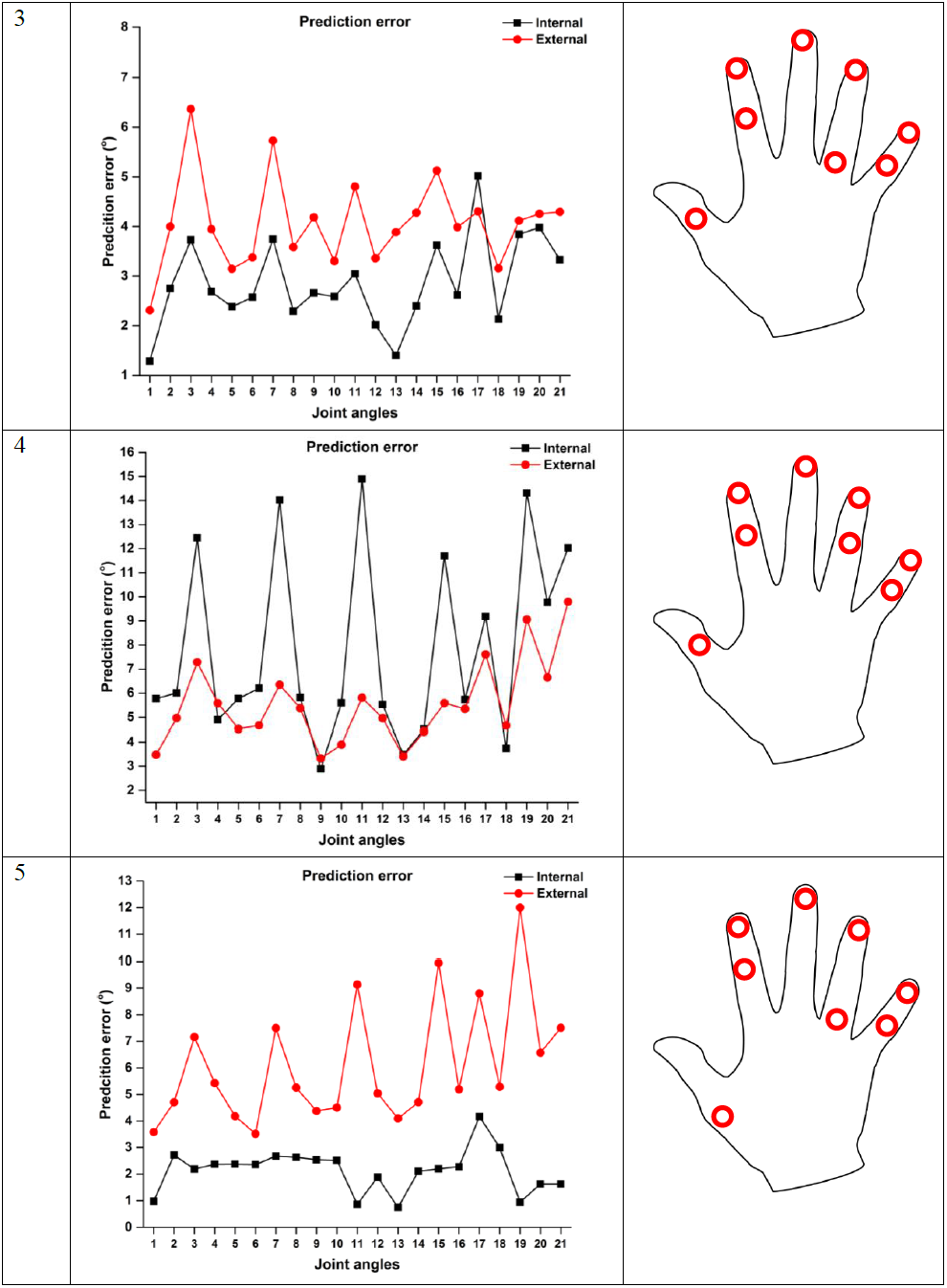
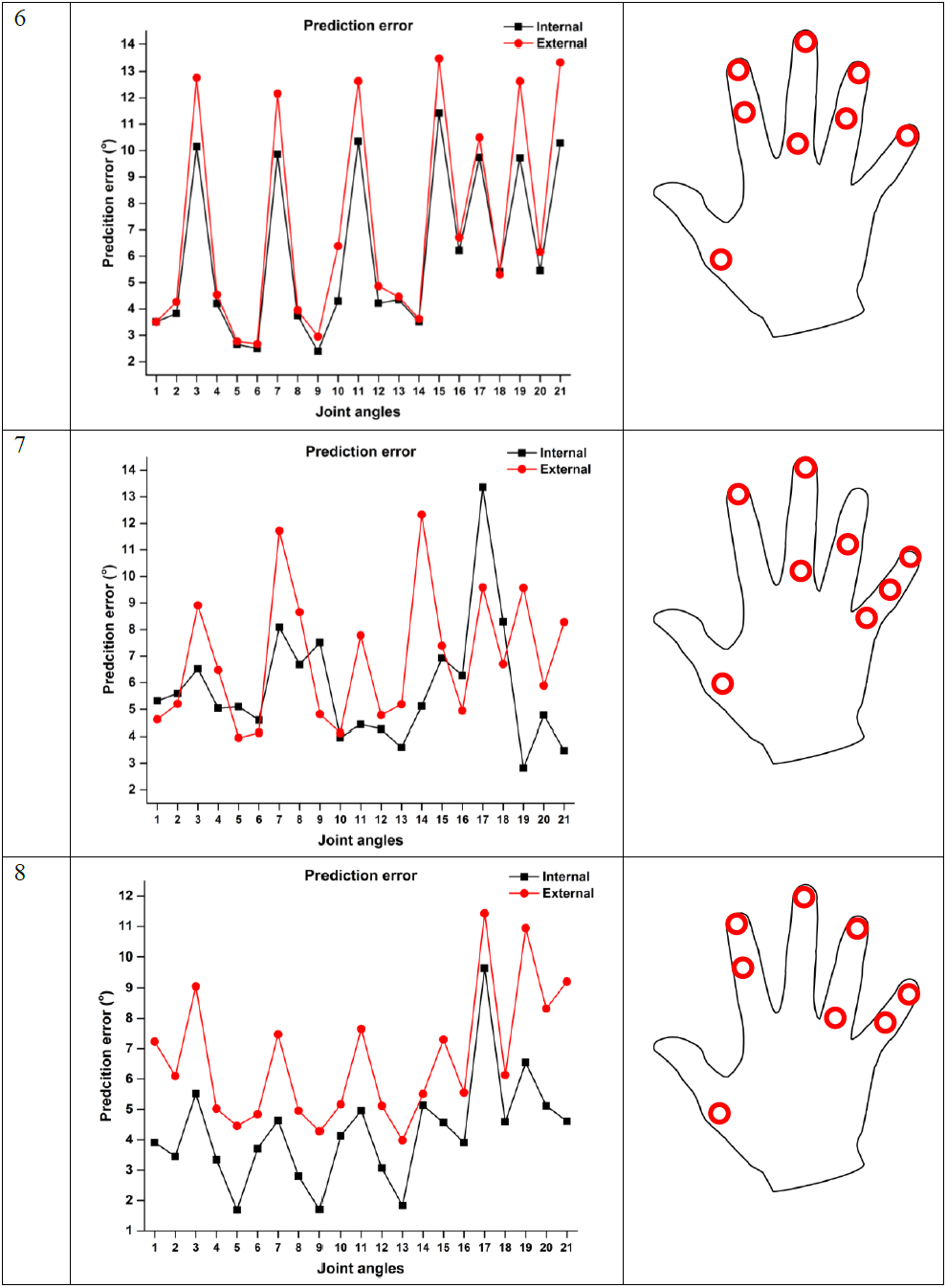
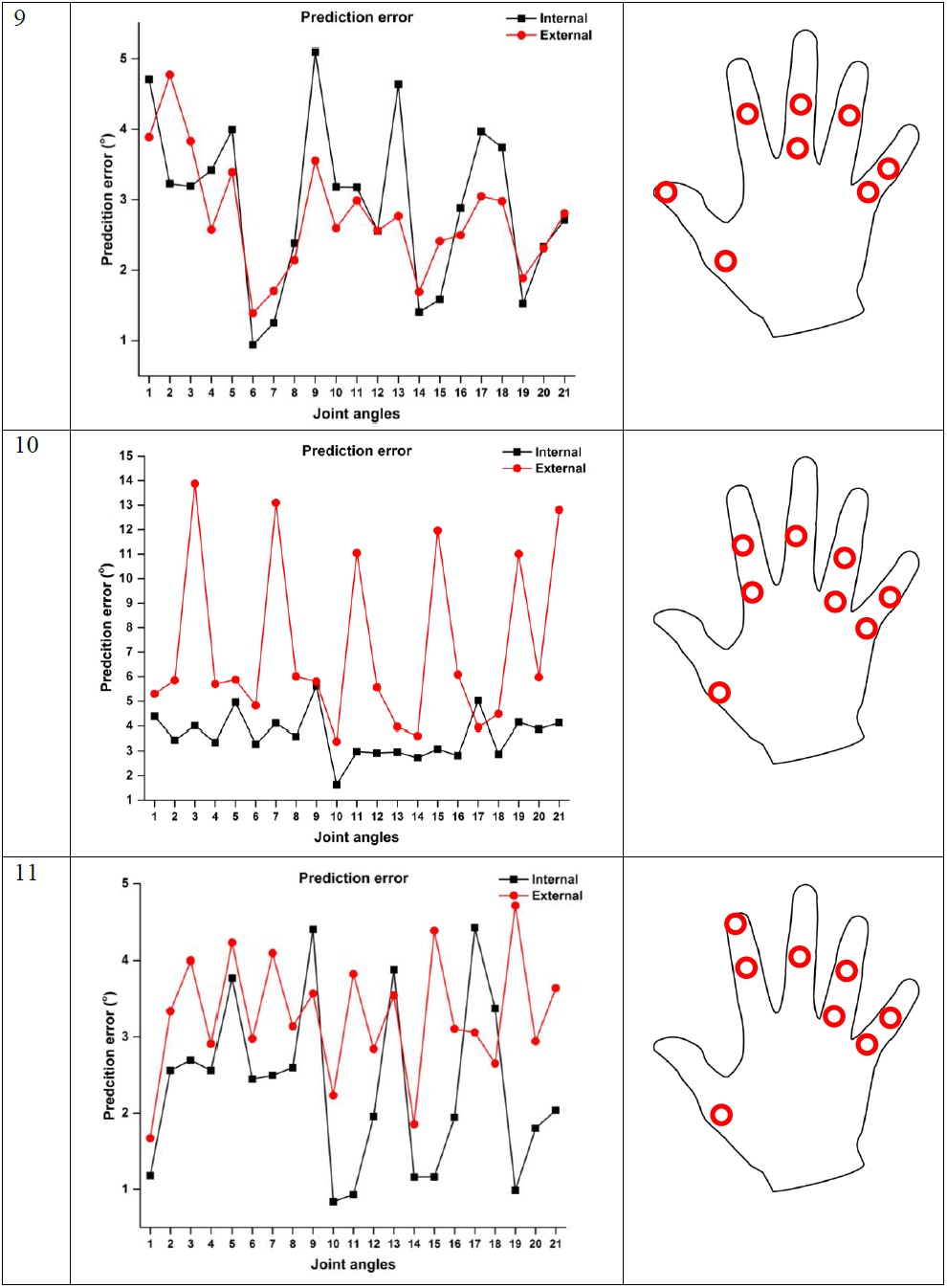
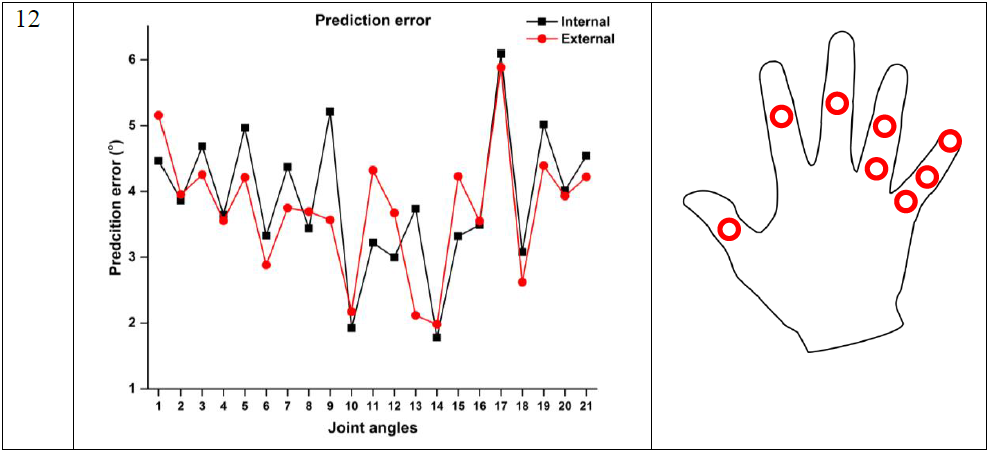
Joint angle reconstruction error for externally constrained postures Vs internally constrained postures.

From the above results we observe that with a single hidden layer, our network model is able to predict the joint angles accurately in the case of internally constrained hand postures. However, the prediction error is large for externally constrained postures for all subjects except for a single subject (subject 4). Joint angle prediction error is smaller for internally constrained postures in comparison with externally constrained postures. Specifically, flexion-extension of MCP joint in all digit shows larger prediction error in comparison other joints in the fingers. ANN with the single hidden layer is not able to capture the flexion-extension movement of MCP joints. Average test error across subjects for externally constrained and internally constrained postures are presented in the figure 20 and 21.

**Figure 20:**
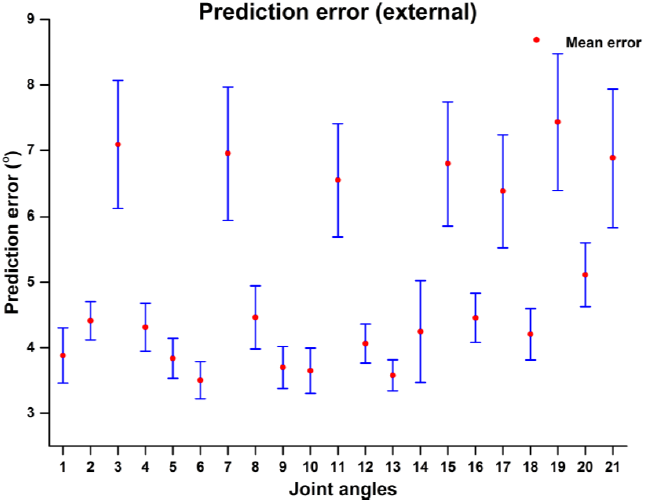
Joint angle prediction error for externally constrained postures average across subjects. Error bar represents standard error of mean across subjects.

**Figure 21:**
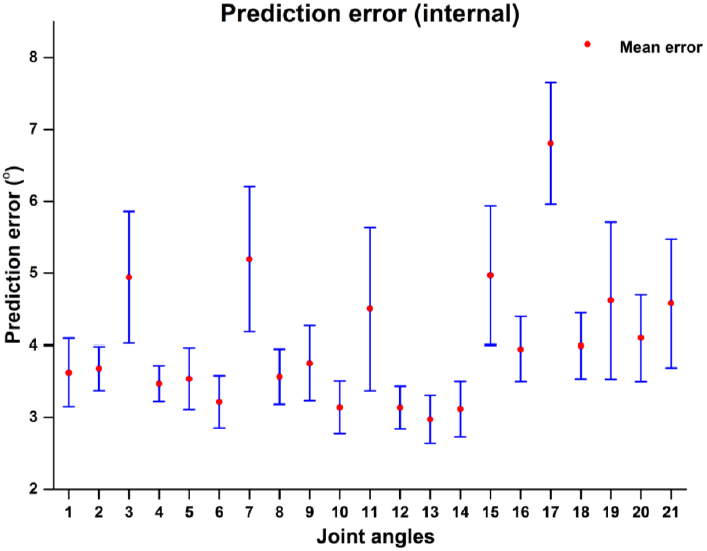
Joint angle prediction error for internally constrained postures average across subjects. Error bar represents standard error of mean across subjects.

Prediction error for externally constrained postures shows large values in flexion-extension movement of MCP joints in comparison with other joint angles. For externally constrained postures mean prediction error across subjects across all joint angles is (5.024 ± 0.47°), whereas for internally constrained postures prediction error is (4.04 ± 0.474°). One of the possible reasons for larger prediction error for externally constrained postures is larger variability in the dataset. In externally constrained postures, subjects cannot specifically grasp the object in the exactly the same way. It might be possible to reduce an error by additional hidden layer in series with the first hidden layer. Joint angles were predicted using two hidden layer network with 90 nodes in the first hidden layer and the second hidden layer consists of 50 nodes. The Levenberg-Marquardt algorithm was used as in the previous case for updating the network weights. The data from selected sensors was used as input to network and output of the network was 21 joint angles. Network performance was measured in terms of mean square error, but for the physical interpretation, root means squared errors are reported in table 6 below.

**Table 6:**
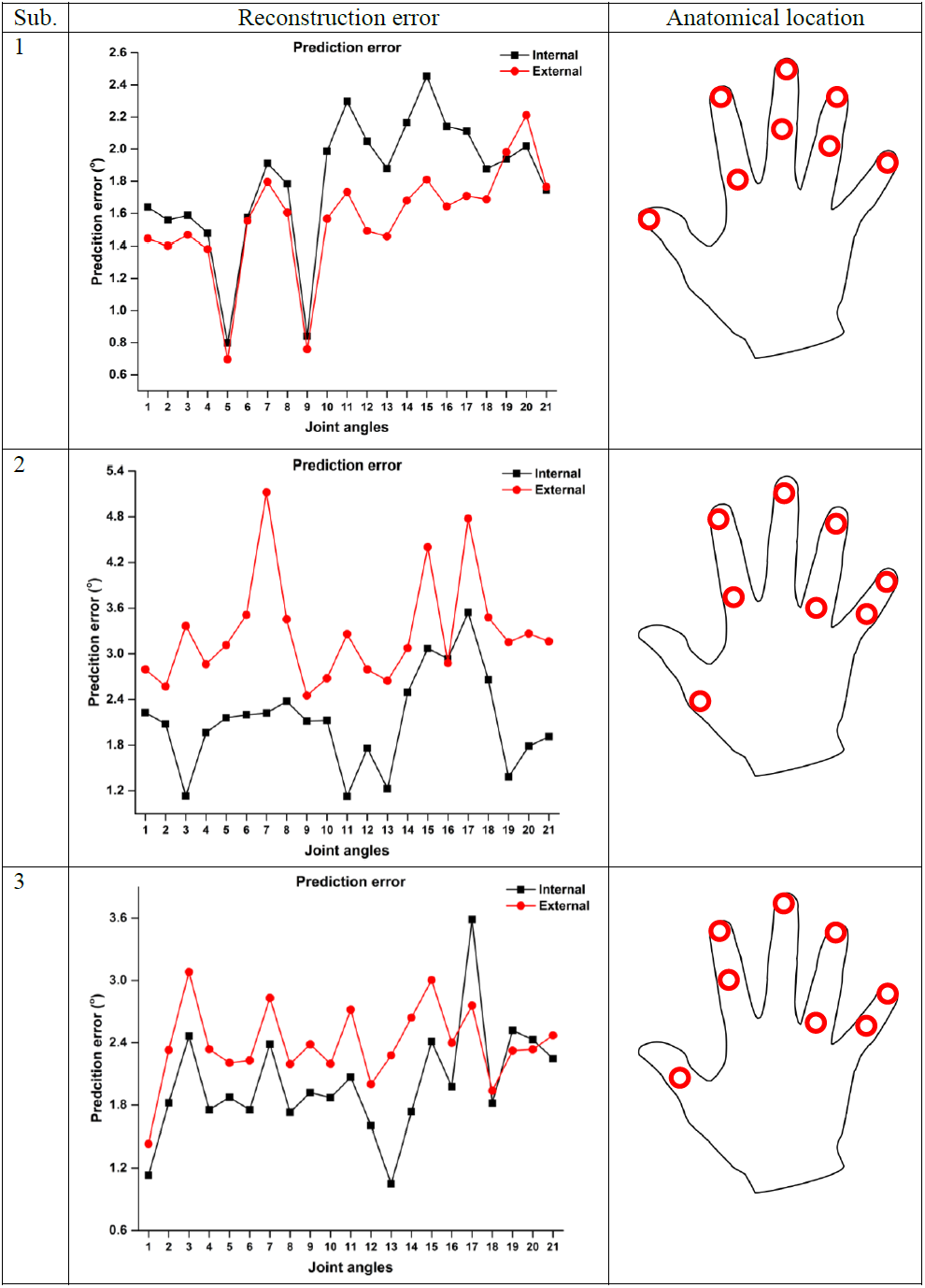
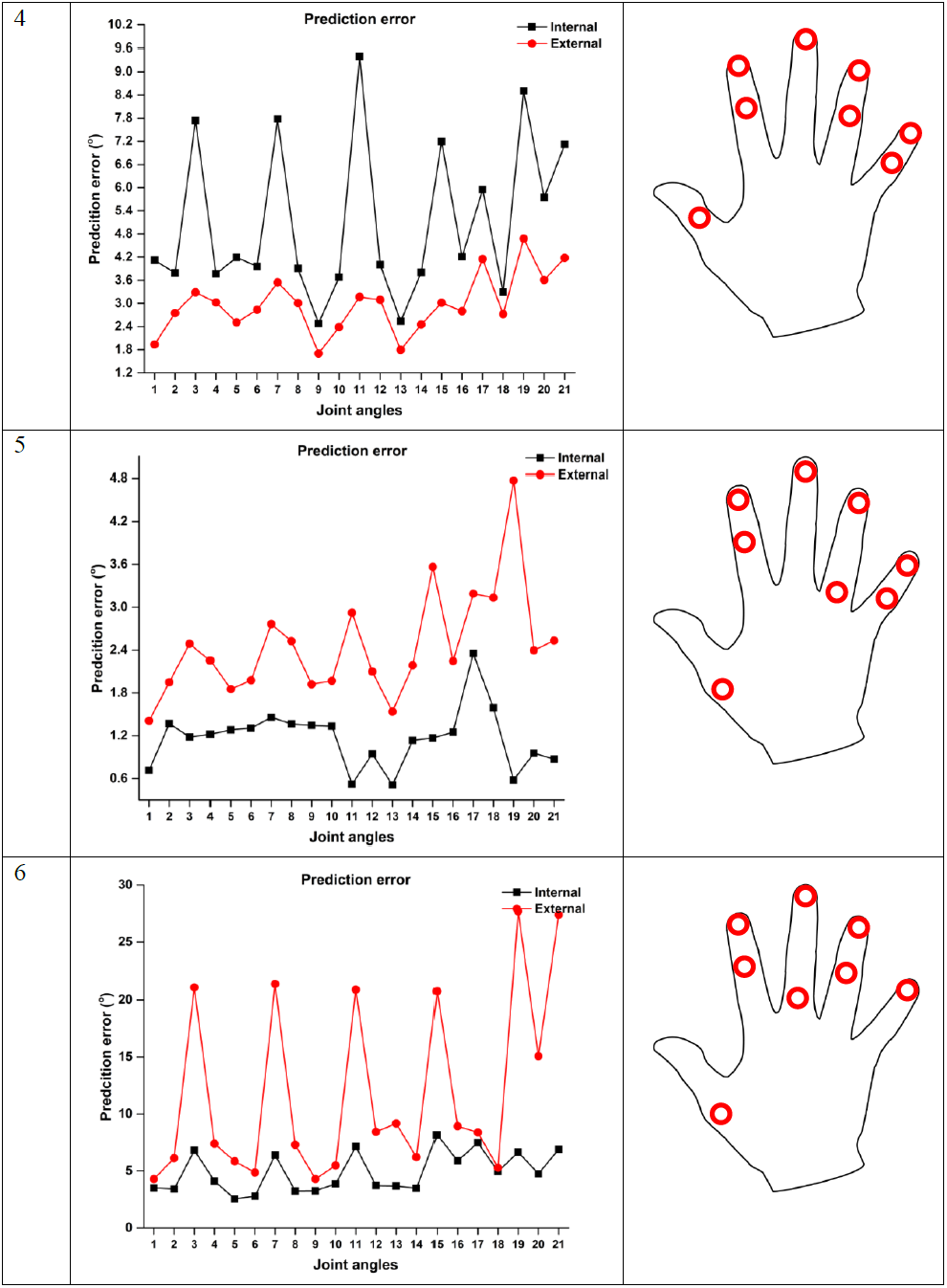
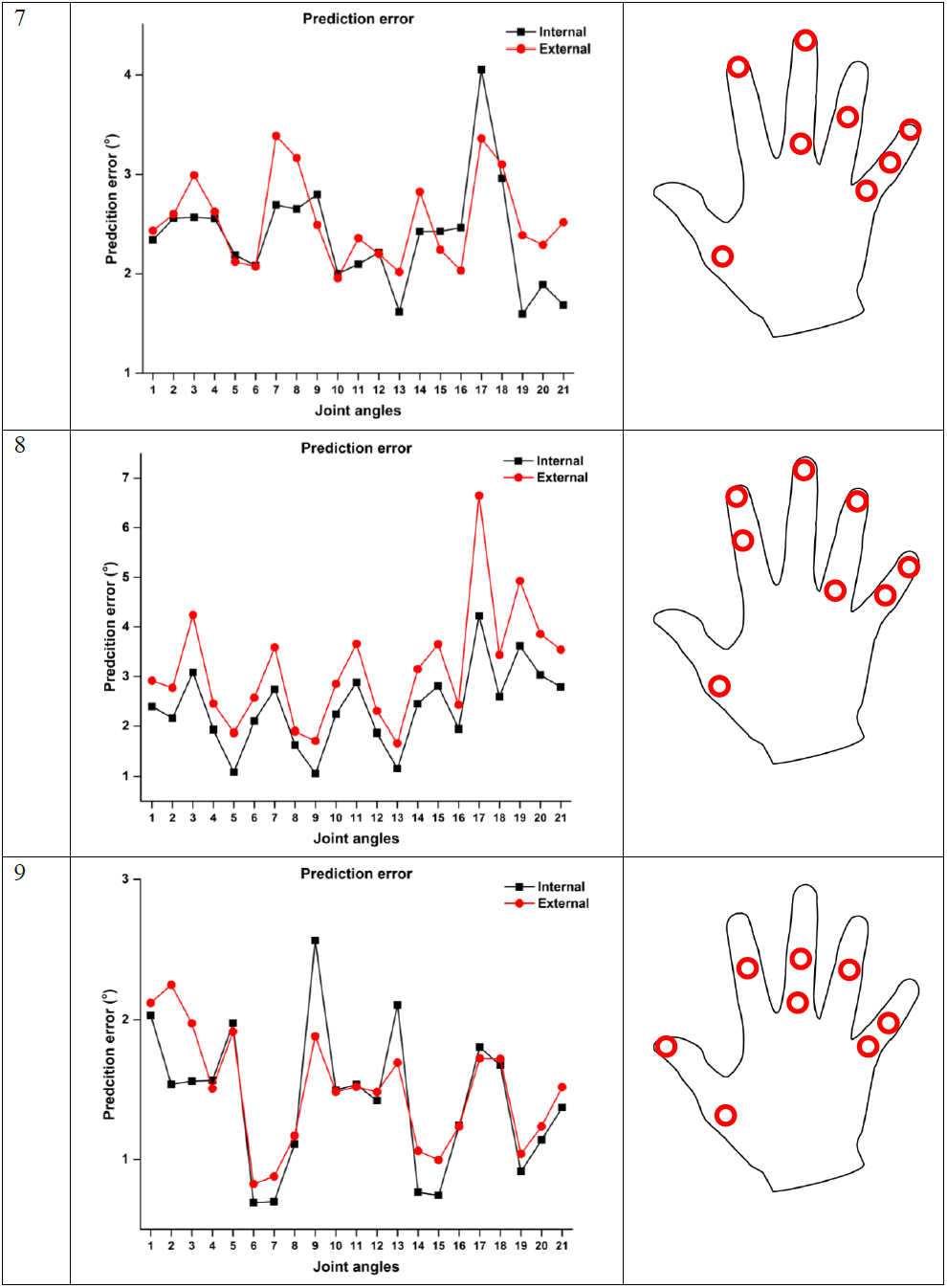
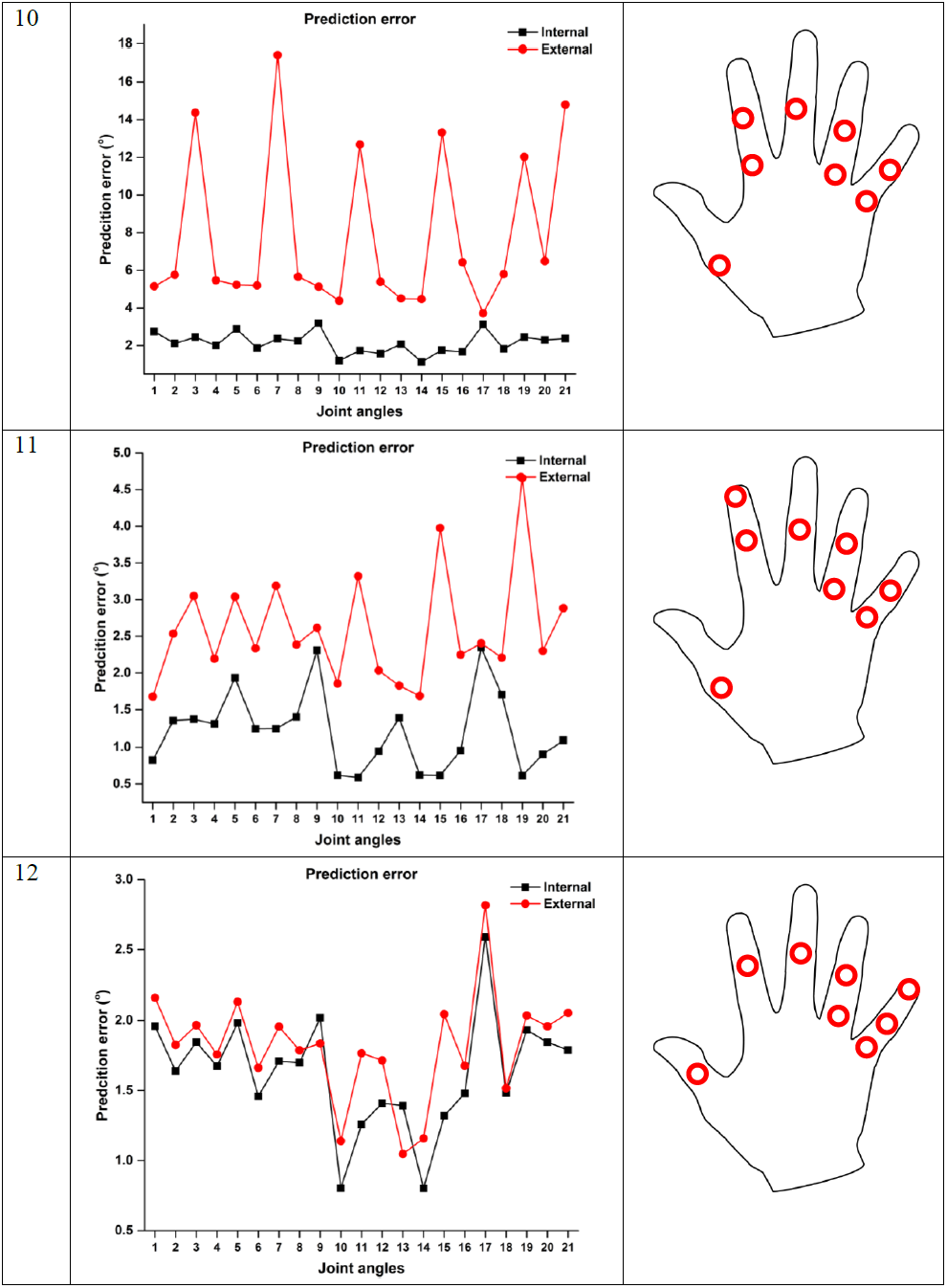
Joint angle reconstruction error for two hidden layers neural network.

**Figure 22:**
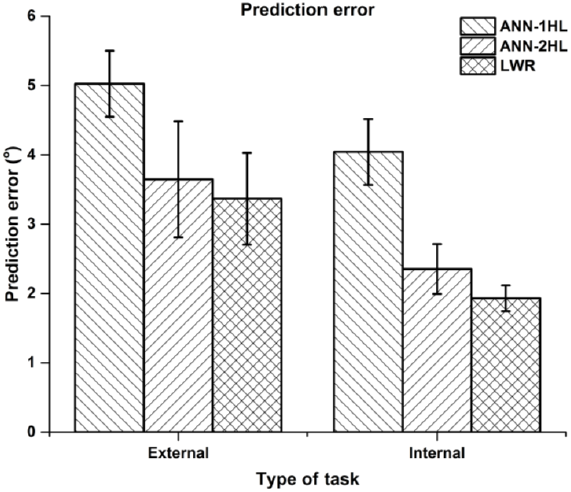
Comparison of prediction error for LWR and ANN with single and two hidden layers. Above result shows larger prediction error in case of an artificial neural network with a single hidden layer in comparison with a network with two hidden layers and locally weighted regression (LWR). We failed to observe any significant difference in prediction of joint angles between the model with 2-hidden layers and LWR. One of the possible reasons for the slightly better performance of LWR is a number of selected sensors are different across subjects. For very few subjects peak of the derivative curve was observed at nearly 2^nd^ or 3^rd^ sensor which demands for a large number of sensors for prediction of joint angles. On the contrary, ANN model consists of ‘8’ sensors for all the subjects. Large prediction error at flexion-extension movement at MCP joint of all the fingers was observed in the case of ANN with 2 hidden layer network similar to ANN with 1 hidden layer case for many subjects. One of the possible reasons for such result could be insufficient training samples for exploring joint angle space for MCP joints.

Our results were compared with previous literature and are presented in table 7 below.

**Table 7:** Comparison of model prediction with previous literature.

| Mora et al. 2012 | Wheatland et al. 2013 | Sensor reduction (our approach) |
| --- | --- | --- |
| **Task:** Reaching and grasping, Pour | **Task:** American Sign Language (ASL) | **Task:** Externally constrained postures, Internally constrained postures |
| Number of Subjects: 6 | Number of Subjects: 1 | Number of Subjects: 12 |
| **Hand model:** 15 DoFs | **Hand model:** 18 DoFs | **Hand model:** 21 DoFs |
| Prediction algorithm: Error backpropagation with ANN | Prediction algorithm: locally weighted regression | Prediction algorithm: Levenberg-Marquardt with ANN |
| **Prediction error:**<br>1HL (100 nodes):<br>**Error:**<br>**$8.64 \pm 1.649^\circ$ (reach and grasp),**<br>**$4.63 \pm 0.828^\circ$ (pour)** | **Prediction error:**<br>**Error:** $22.9^\circ$ using 6 markers for ASL (i.e. subset of internally constrained postures) | **Prediction error:**<br>**Error:** **$(3.3657 \pm 0.66)^\circ$** for externally constrained postures and **$(1.9315 \pm 0.187)^\circ$** for internally constrained postures using LWR model. Selected sensors changes for different subjects. |
| **Prediction error:**<br>2HL (150,100 nodes):<br>**Error:**<br>**$15.12 \pm 2.74^\circ$ (reach and grasp),**<br>**$14.95 \pm 5.88^\circ$ (pour)** |  | **Prediction error:**<br>1HL (30 nodes)<br>**Error:**<br>**$(4.0411 \pm 0.474)^\circ$ for internally constrained postures,**<br>**$(5.0245 \pm 0.475)^\circ$ for externally constrained postures using 8 sensors** |
|  |  | **Prediction error:**<br>2HL (90,50 nodes):<br>**Error:**<br>**$(2.353 \pm 0.359)^\circ$ for internally constrained postures,**<br>**$(3.64 \pm 0.837)^\circ$ for externally constrained** |

The comparative table is presented above for two different models (LWR and ANN). Hand model chosen is of 21 DoF in comparison with previous work. Prediction error in all cases is smaller for LWR and ANN model compared to previous studies. Our systematic sensor selection approach and model for prediction of joint angle shows promising results in comparison with previous studies.

### 1.7 Discussion

#### 1.7.1 Preliminary sensor selection based on PCA

From the PCA based sensor ranking algorithm, sensors located near distal part of a finger are critical in comparison with sensors mounted on other locations. One of the possible reasons for these results is that for various hand postures, the distal part has larger variance and contributes more in space. Hence, collecting kinematic information from the distal part of the hand may help in exploring larger action manifold of hand. PCA sensor ranking algorithm shows higher rank (i.e. more importance) for sensors mounted on distal phalanges of a particular finger. These sensors may provide critical information about hand postures and hand movements. Sensor selection is subject specific. Thumb sensor mounted on distal and proximal phalange (i.e. sensor 13 and 15) is important for generating stable grasp while performing a task involving activities of daily living (ADL). Capturing thumb movements is necessary to describe the opposition movement of thumb (Napier 1956; 1993). The mobility of thumb is much larger than any other digit in hand and hence more sensors are required for modeling thumb position accurately. Previous literature observed that for classification of different grasps, the position of thumb is not important, hence monitoring thumb position is not needed (Chang et al., 2007). Our finding does not support these results. Our findings show that for accurate hand posture reconstruction thumb sensors are important.

#### 1.7.2 PCA ranking algorithm shows evidence for finger co-variation

PCA ranking algorithm allows determining important sensors based on variance covered by a sensor in orientation space. Ranking the sensors will allow us to generate *sensor importance curve*. Sensor importance (S.I.) curve was computed for each subject separately. Correlation analysis was performed on S.I. curve and correlation matrix was derived. This matrix is presented in figure 23 below. This result shows that sensors mounted on middle and ring are highly correlated. This result implies that for all the subjects, middle and ring finger sensors show high correlation values.

**Figure 23:**
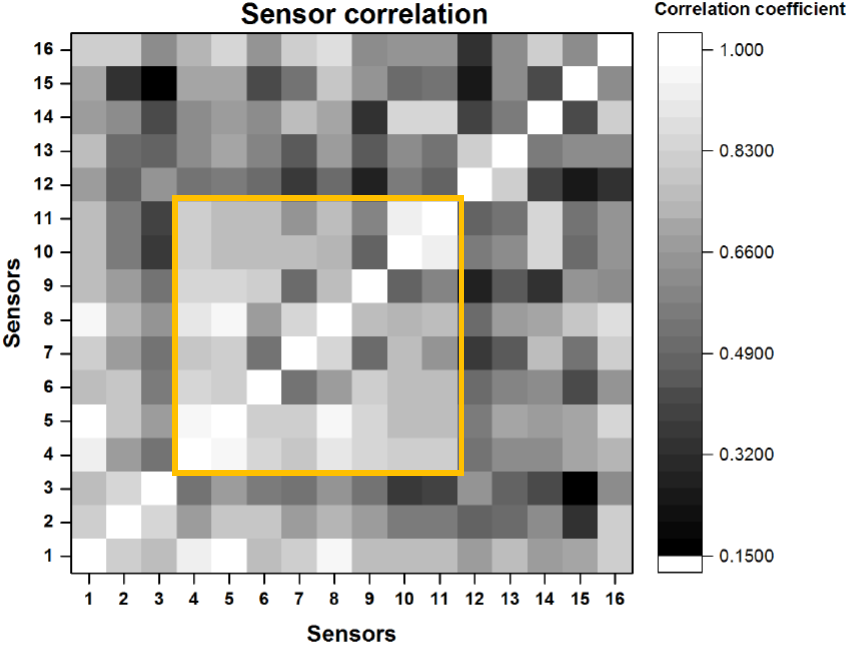
Correlation coefficient between sensors. Checkerboard plot shows a correlation between all sixteen sensors based on the contribution of each sensor based on PCA ranking algorithm for all twelve subjects. The marked square region shows a higher correlation between sensors. Rank of these sensors is from 4-11 which are mounted on middle, ring and little fingers showing a high correlation between sensors.

One of the possible reasons for large correlation between middle and ring finger sensors is covariation between these fingers. This is also called as *enslavement effect* (Ross and Schieber 2000). Our result shows strong evidence for middle and ring finger co-variation in all subjects.

#### 1.7.3 Sensor correlation will help in determining redundant sensors

Strong linear relationship was observed between distal interphalangeal joint (DIP) and proximal interphalangeal joint (PIP) across subjects. One can argue that by measuring one of these joint movements, it is possible to calculate the movement of the other joint. It is possible to reduce the number of sensors by finding a correlation between sensors that allows reconstruction of one sensor value based on the value of a correlated sensor. In this chapter, a method for determining a correlation between sensors with multiple outputs (i.e. azimuth, elevation, and roll) is presented. Finding correlation between sensors in each orientation separately helps us finding equivalent sensors for a given orientation. The threshold value for determining correlation allows physical relevance for a given orientations. Combining all the orientation correlations gives a possible list of redundant sensors. Our findings suggest a strong correlation between sensor mounted on distal and middle segments across subjects. Simultaneous flexion movement of DIP and PIP joints are controlled possibly due to anatomical structure and sharing of muscle and tendons (i.e. Flexor digitorum profundus) which cause flexion movements (Napier and Tuttle 1993). Sensor correlation is not mere by-product of our mathematical formulation but there is physical and physiological significance to it.

#### 1.7.4 Network performance based sensor ranking

For addressing the hidden nonlinearity between joint angles, neural network is a potential candidate. Hence degradation in performance (i.e. ability to predict joint angles) due to an elimination of a sensor can be used for ranking the sensors. If elimination of a sensor generates a large mean squared error in performance, that particular sensor is important. Due to the structural difference on a subject basis, subject specific models were obtained and subject specific ranking was obtained.

#### 1.7.5 Systematic approach for sensor selection

In many cases, reducing markers for tracking hand movements is performed using an iterative method (Hoyet et al., 2012) or based on PCA ranking algorithm (Wheatland et al., 2013). Few studies showed removing markers from middle and ring fingers and monitoring index and little fingers are enough to reconstruct hand posture from a perceptual point of view. This result does not account for finger co-variations and enslavement effects present in human hand. Hence, to overcome this issue a systematic approach is proposed which accounts for action manifold spanned by all five fingers, the correlation between sensors and network performance. The method gives a possible set of sensors from the data-driven model and mechanics perspective. The method is flexible for selecting a number of sensors based on requirement. In our case, we reduced the number of sensors from 16 to 8.

#### 1.7.6 Joint angle prediction using Locally Weighted Regression

Locally weighted regression is a nonparametric based model for joint angle reconstruction of all 21 joints from a reduced set of sensors. Movements and hand postures are spread in high dimensional space. We used PCA ranking algorithm for selecting reduced set of sensors and prediction of 21 joint angles (Wheatland et. al., 2013).

LWR shows relatively small prediction error in DIP and MCP joints of all four fingers and relatively large prediction error in PIP joints of middle, ring and little fingers. Largest prediction error was observed for PIP joint of the ring finger (i.e. joint 10) in all cases across subjects. One possible reason for large prediction error is a large degree of dependency in middle and ring fingers in comparison with index and little fingers (Li et al. 2004). The study exhibit index and little finger shows larger degrees of functional independence during the performance of activities of daily living. During flexion movement of fingers, low-threshold motor units acting on neighboring digits are recruited in flexor digitorum profundus (FDP), flexor digitorum superficialis (FDS) and extensor digitorium (ED) muscles (Kilbreath and Gandevia 1994; Butler et al. 2005; van Duinen et al. 2009). Our data-driven model is limited to accurate prediction of independent fingers. However model cannot capture enslavement effects. This might result in larger prediction error of PIP joints. Sensor correlation in middle and ring finger provide support to this argument.

Our observations show that locally weighted regression model is able to capture and predict joint angles for internally constrained hand postures accurately. However, the model fails to predict joint angles for externally constrained hand postures. There are two possible reasons for the above result. One, large variability in the externally constrained task. Participant uses a different strategy for achieving the task objective. Due to multiple degrees of freedom in hand, subject tries to use a different strategy for achieving the goal of grasping an object. Second, locally weighted regression is a nonparametric approach. If the number of samples is small, joint angle prediction error could be large. In addition to that if the distribution of dataset is relatively sparse then it is difficult to model using distance-based approach.

Using LWR, prediction error for **internally constrained hand postures** is **(1.9315 ± 0.187°)** and for **externally constrained hand postures** is **(3.3657 ± 0.66°)**. Previously we observed stereotypical pattern for achieving task successfully. This pattern was relatively consistent across subjects. Hence we can observe lesser variability for internally constrained postures. In comparison with, externally constrained hand postures, there are more internally constrained hand postures which may contribute for predicting joint angles accurately. A major drawback of this approach is computation time for prediction of a query. As a number of samples increases, the model needs to allocate weights based on nearest neighbour points. With lesser number of samples, the model is unable to predict joint angle accurately. Hence joint angle prediction needs an approach for predicting joint angles relatively small prediction error and less time-consuming.

#### 1.7.7 Joint angle prediction using artificial neural network

Artificial neural network with single hidden layer and two hidden layers were used for prediction of joint angles from a reduced set of sensors. The advantage of this approach over locally weighted regression is that once the network is trained we are able to predict joint angle with significantly lesser time in comparison with the locally weighted regression. Joint angle prediction error is larger in externally constrained hand posture in comparison with internally constrained hand postures. One of the possible reasons, is that a large number of training samples are available in comparison with internally constrained postures. Another possible reason is finger covariation during externally constrained postures which may introduce *enslavement effect* in middle and ring finger and neural network model is not able to account for it.

ANN model with a **single hidden layer (30 nodes)** generates prediction error of **(4.041 ± 0.47°) for internally constrained postures** and **(5.024 ± 0.475°) for externally constrained postures**. Average joint angle prediction error was observed to be larger in externally constrained postures in comparison with internally constrained hand postures. The result shows ANN with two hidden layers predicts hand postures accurately with significantly less computational time compared to LWR method. The advantage of this approach is a number of sensors are *8* for all the subjects unlike in LWR. The combination of a systematic approach to reduce the number of sensors and joint angle prediction model allows for faithful reproduction of hand movements with less computation time and less complexity. In table 4-8 comparison of proposed approach with previous literature is presented. Joint angle prediction with two hidden layers ANN shows promising results with 8 sensors for the more complex model (21 DoFs). Results in the previous literature use 15 DoF model with more complex ANN (150,100) nodes for six subjects. Mean prediction error of (15.24 ± 2.74**°**) and (14.95 ± 5.88**°**) for reach-to-grasp and pour task respectively were reported. The method we proposed uses the network with **2 hidden layers** with **(90, 50) nodes** respectively. Mean prediction error for **internally constrained postures** is **(2.353 ± 0.359°)** and for **externally constrained postures** is **(3.64 ± 0.837°)**. We believe that our method performs better than previous methods because in our method, sensors are selected based on a systematic approach rather than arbitrarily.

This chapter proposed a systematic approach for selecting a set of sensors based on the user requirement and network performance. Our method is flexible enough to account for a different set of sensors which can predict joint angles accurately. Hence, a combination of proposed method and neural network model will help in reproducing finger movements accurately. This will help prediction of hand postures for the field for robotics, biomechanics and animation industry.

